# PIEZO1 overexpression in hereditary hemorrhagic telangiectasia arteriovenous malformations

**DOI:** 10.1101/2024.11.27.625696

**Authors:** Hyojin Park, Sungwoon Lee, Jessica Furtado, Mark Robinson, Martin Schwartz, Lawrence Young, Anne Eichmann

**Affiliations:** Cardiovascular Research Center, Department of Internal Medicine, Yale University School of Medicine, New Haven CT, USA; Yale University School of Medicine, Department of Molecular and Cellular Physiology; Yale University School of Medicine, Department of Hematology; Yale University School of Medicine, Departments of Cell Biology and Biomedical Engineering; Université de Paris, PARCC, INSERM, F-75006 Paris

## Abstract

**Background:** Hereditary hemorrhagic telangiectasia (HHT) is an inherited vascular disorder characterized by arteriovenous malformations (AVMs). Loss-of-function mutations in Activin receptor-like kinase 1 (ALK1) cause type 2 HHT and *Alk1* knockout (KO) mice develop AVMs due to overactivation of VEGFR2/PI3K/AKT signaling pathways. However, the full spectrum of signaling alterations in *Alk1* mutants remains unknown and means to combat AVM formation in patients are yet to be developed.

**Methods:** Single-cell RNA sequencing of endothelial-specific *Alk1* KO mouse retinas and controls identified a cluster of endothelial cells (ECs) that was unique to *Alk1* mutants and that overexpressed fluid shear stress (FSS) signaling signatures including upregulation of the mechanosensitive ion channel PIEZO1. PIEZO1 overexpression was confirmed in human HHT lesions, and genetic and pharmacological PIEZO1 inhibition was tested in *Alk1* KO mice, as well as downstream PIEZO1 signaling.

**Results:** Pharmacological PIEZO1 inhibition, and genetic *Piezo1* deletion in *Alk1*-deficient mice effectively mitigated AVM formation. Furthermore, we identified that elevated VEGFR2/AKT, ERK5-p62-KLF4, hypoxia and proliferation signaling were significantly reduced in *Alk1*-*Piezo1* double ECKO mice.

**Conclusions:** PIEZO1 overexpression and signaling is integral to HHT2, and PIEZO1 blockade reduces AVM formation and alleviates cellular and molecular hallmarks of ALK1-deficient cells. This finding provides new insights into the mechanistic underpinnings of ALK1-related vascular diseases and identifies potential therapeutic targets to prevent AVMs.

## Introduction

Hereditary hemorrhagic telangiectasia (HHT) is a rare inherited autosomal dominant vascular disorder that affects approximately 1 to 2 per 10,000 individuals worldwide^1, 2^. HHT patients develop arteriovenous malformations (AVMs), which are abnormal direct connections between arteries and veins that bypass capillaries. Small AVMs in the skin and mucous membranes are referred to as telangiectasias; their rupture can lead to frequent epistaxis, GI bleeding, and anemia, which significantly impacting the quality of life for those with HHT^2, 3^. Larger AVMs in the liver, lungs, or brain may cause severe, life-threatening conditions such as high-output heart failure and stroke^4^.

Over 90% of HHT cases are linked to heterozygous mutations in the endothelial surface receptors ENG (Endoglin, mutated in HHT1) and ALK1 (ACVRL1, mutated in HHT2)^5-8^. Additionally, mutations in SMAD4 are responsible for a combined juvenile polyposis–HHT syndrome, accounting for less than 5% of HHT cases^9^. ALK1 and ENG are receptors for the transforming growth factor–β (TGF-β) superfamily members BMP9 and BMP10. Ligand binding activates ALK1/ENG receptor signaling, which triggers the phosphorylation of cytoplasmic SMAD1/5/8 proteins and complex formation with SMAD4, leading to nuclear translocation and regulation of a set of genes that prevent AVM formation. Mutations associated with HHT disrupt various components of this endothelial signaling pathway.

Despite extensive study of HHT genetics, the underlying cellular and molecular mechanisms, including the pathways altered by loss of ALK1 signaling, remain unclear, hindering the development of new treatment options. Recent findings by others and us revealed that AVM lesions originate from loss of HHT signaling in venous and capillary endothelial cells (ECs). This loss leads to altered polarization, migration against the direction of blood flow, increases in EC size, accelerated proliferation, and defective arterio-venous gene expression that contribute to AVM formation^10-12^. All changes occur in response to fluid shear stress (FSS), and arresting blood flow in zebrafish HHT mutants rescues AVM formation^5, 13^.

Mechanistically, blocking endothelial BMP9-ALK1-ENG signaling leads to increased activation of pro-angiogenic vascular endothelial growth factor receptor 2 (VEGFR2) signaling at the plasma membrane, and overactivation of intracellular phosphoinositide 3-kinase (PI3K) and AKT^10, 14-16^. Pharmacologic inhibition of VEGFR2 or PI3K prevents AVM formation in *Alk1*-deficient mice and reduces the diameter of AVMs in *Eng* mutants^14, 17^. Elevated PI3K signaling has been confirmed in biopsies of cutaneous telangiectasias from patients with HHT type 2, and VEGFR2 inhibition is approved for treatment of HHT patients^16, 18^. However, VEGFR2 inhibition shows variable efficacy and does not universally prevent AVM progression, suggesting that targeting this pathway alone may not fully address the complexity of AVM pathogenesis. Furthermore, while elevated PI3K signaling is implicated in AVM formation, overactivation of PI3K alone is insufficient to induce AVMs, indicating that additional signaling alterations are necessary to drive the disease.

To identify the spectrum of signaling alterations in AVMs, we performed single cell (sc)-RNA sequencing analysis of *Alk1* mutant ECs, which identified a unique cluster of ECs with altered arterio-venous identity. Among the upregulated pathways in this cluster, we identified increased PIEZO1-driven FSS signaling^19, 20^. PIEZO1 is an endothelial mechanosensitive ion channel known to mediate the endothelial response to FSS^21-24^, prompting us to investigate its contribution to HHT signaling. We demonstrate that PIEZO1 is overexpressed in human HHT lesions and functions as an upstream regulator of aberrant EC behavior in *Alk1* mouse mutants and in human ECs *in vitro*. Our findings identify new targets to prevent vascular malformations in HHT patients with ALK1 mutations.

## Materials and methods

### Mice

All animal experiments were performed under a protocol approved by the Institutional Animal Care Use Committee of Yale University. *Alk1^f/f^* mice were kindly provided by Dr. S. Paul Oh^25^. *Cdh5 Cre^ERT2^* mice were kindly provided by Dr. Ralf Adams^26^ and *Mfsd2a Cre^ERT2^* mice were kindly provided by Dr. Bin Zhou^27^. *Piezo1^f/f^* mice were purchased from the Jackson Laboratory (Strain #:029213). Seven to eight weeks old *Alk1^f/f^*, *Piezo1^f/f^*, *Cdh5 Cre^ERT2^* or *Mfsd2a Cre^ERT2^* mixed genetic background were intercrossed for experiments and *Alk1^f/f^ Cdh5 Cre^ERT2^*, *Alk1^f/f^ Mfsd2a Cre^ERT2^*, *Alk1^f/f^ Piezo1^f/f (or f/+)^ Cdh5 Cre^ERT2,^ Alk1^f/f^ Piezo1^f/f (or f/+)^ Mfsd2a Cre^ERT2^* mice were used. Gene deletion was induced by intra-gastric injections with 100 μg Tamoxifen (TAM) (Sigma, T5648; 2.5 mg ml^−1^) into pups at P4. TAM-injected *Cre^ERT2^* negative littermates were used as controls.

### Latex dye injection

P6 pups were anesthetized on ice, and abdominal and thoracic cavities were opened. The right atrium was cut, blood was washed out with 2 ml PBS and 1 ml of latex dye (VWR, LL0057-1L) was slowly and steadily injected into the left ventricle with an insulin syringe. Retinas and GI tracts were washed in PBS and fixed with 4% paraformaldehyde (PFA) overnight. Brains and GI tracts were cleared in Benzyl Alcohol: Benzyl Benzoate (1:2) for 2-3 days before imaging.

### Reagents and antibodies

For immunostaining: IB4 ([IsolectinB4] #121412, 1:500; Life Technologies), Alexa Fluor 488 (#A-21311, 1:1000; Invitrogen), mouse anti-ALK1 (#AF770, 1:300; R&D) human anti-ALK1(#AF370, 1:300; R&D), DAPI (#D1306, 1:1000; Life Technologies), mouse KLF4 (#AF3158, 1:300; R&D), human KLF4 (#AF3640, 1:300; R&D), PIEZO1(#15939-1-AP, 1:200, Proteintech), SQSTM1/p62 (ab109012, 1:300, Abcam), Mouse KLF4 (#AF3158,1:200, R&D), Human KLF4 (#AF3640, 1:200, R&D), ERK5 (#E1523-.2ML,1:200, Sigma), CD31 (#AF3628, 1:200, R&D), Alexa Fluor 647-Anti-ERG (AB196149, 1:500, abcam) For western blotting: anti-ALK1 (7R-49334, 1:1000; Fitzgerald), β-actin (#A1978 1:3000; Sigma), ERK5 (#E1523-.2ML,1:2000, Sigma), Phospho-ERK5 (#3371, 1:1000, Cell Signaling Technology), PIEZO1 (NBP1-78537,1:1000, Novus Biologicals) Phosho-AKT(#4060,1:1000, Cell Signaling Technology), AKT (#4691,1:1000, Cell Signaling Technology), KLF4 (ab125036,1:1000, Abcam) Phospho-VEGFR2 (#2478, 1:1000, Cell Signaling Technology) VEGFR2 (#9698,1:1000, Cell Signaling Technology) HIF1A (#14179, 1:1000, Cell Signaling Technology) Appropriate secondary antibodies were fluorescently labeled (Alexa Fluor donkey anti-rabbit, Alexa Fluor donkey anti-goat) or conjugated to horseradish peroxidase (anti-rabbit and anti-mouse IgG [H+L], 1:8.000; Vector Laboratories).

GsMTx4 (P1205, Selleckchem, 1 mg/kg) was administered intraperitoneally (i.p.) on postnatal days (P) 4 and 5 following TAM injection into the P4 mouse.

### Immunostaining

The eyes of P6/P8 pups were prefixed in 4% formaldehyde for 8 minutes (min) at room temperature. Retinas were dissected, blocked for 30 min at room temperature in blocking buffer (1% fetal bovine serum, 3% BSA, 0.5% Triton X-100, 0.01% Na deoxycholate, 0.02% Sodium Azide in PBS at pH 7.4) and then incubated with specific antibodies in blocking buffer overnight at 4°C. The next day, retinas were washed and incubated with IB4 together with the corresponding secondary antibody overnight at 4°C. The next day, retinas were washed and post-fixed with 0.1% formaldehyde and mounted in fluorescent mounting medium (DAKO, USA). High-resolution pictures were acquired using ZEISS LSM800 and Leica SP8 confocal microscope with a Leica spectral detection system (Leica TCS SP8 detector), and the Leica application suite advanced fluorescence software. Quantification of retinal vasculature was done using ImageJ and then Prism 7 software for statistical analysis.

For cell immunostaining, cells were plated on gelatin-coated dishes. Growing cells were fixed for 10 min with 4% PFA and permeabilized with 0.1% Triton X-100 for 10 min prior to overnight incubation with primary antibody and then secondary antibody conjugated with fluorophore.

### Detection of hypoxia

Hypoxia was detected by injecting 60 mg/kg pimonidazole 90 min before euthanasia. Retinas were then stained using the Hypoxyprobe Kit (#HP1-200Kit, Hypoxyprobe) with Hypoxyprobe Mab1-FITC (#FITC-Mab, 1:100, Hypoxyprobe).

### Cell culture and siRNA transfection

Human umbilical vein endothelial cells (HUVECs) were purchased from Lonza (C2519A) and cultured in EGM2-Bullet kit medium (CC-3156 & CC-4176, Lonza). Depletion of *ALK1* or *PIEZO1* was achieved by transfecting 20 pmol of small interfering RNA (siRNA) against *ALK1* (Qiagen, mixture of 2 siRNAs: S102659972 and S102758392) or *PIEZO1* (Invitrogen, mixture of 2 siRNAs: S18893 and S18892) using Lipofectamine RNAiMax (Invitrogen). Transfection efficiency was assessed by western blotting and quantitative PCR (qPCR). Experiments were performed 72 hr post transfection and results were compared with siRNA CTRL (ON-TARGETplus Non-Targeting Pool D-001810-10-05).

### Exposure of Endothelial Cells to Shear Stress

Confluent HUVEC monolayers were grown in 6-well plates. Sixty hr after CTRL, *ALK1* and/or *PIEZO1* siRNA transfection, cells were exposed to 45 min-16 hr shear stress, using an orbital shaker to generate laminar shear stress of ∼10 dynes/cm^2^.

### Western blotting

Cells were lysed with Laemmli buffer including phosphatase and protease inhibitors (Thermo Scientific, 78420, 1862209). 20 μg of proteins were separated on 4% to 15% Criterion precast gels (567–1084, Biorad) and transferred on 0.23 um nitrocellulose membranes (Biorad). Western blots were developed with chemiluminescence horseradish peroxidase substrate (Thermo Scientific #34096) on a Luminescent image Analyzer, Gel Doc XR+ Documentation System (Biorad). Bands were quantified using ImageJ.

### Proliferation Assay

Proliferation analysis was conducted using the Click-iT EdU Alexa Fluor 488 Imaging Kit (Life Technologies). P6 pups were injected with 200 μg of EdU (5 mg/mL) and euthanized 4 hrlater. EdU staining was performed according to the manufacturer’s protocol.

### Quantitative Real-Time PCR primers

Mm_Acvrl1 (Qiagen, QT00161434), Mm_Gapdh (Qiagen, QT01658692), Hs_ACVRL1 (Qiagen, QT00050351), Hs_GAPDH (Qiagen, QT00079247), Mn_Piezo1 (FW:CATTA TCCTCTTCCTCATCGCC, REV:ATGGTGAACAGTGGCTCATAG), Hs_PIEZO1 (FW:C AATGAGGAGGCCGACTACC, REV:GCACTCCTGCAGTTCGATGA), Mm_Klf4 (FW:GA GTTCCTCACCGGAACG, REV:CGGGAAGGGAGAAGACACT) Quantitative real-time PCR was performed in a real-time PCR system (BioRad CFX96 Real Time PCR Thermal Cycler)

### sc-RNA seq Library preparation, sequencing and bioinformatics analysis

Single cell suspensions of retinal cells were resuspended in PBS containing 0.04% ultra-pure BSA^28^. ScRNA-seq libraries were prepared using the 10x Genomics Chromium Next GEM Single Cell 3′ Library Construction Kit V3.1 (CG000204. Generated libraries were sequenced on an Illumina HiSeq4000 for 150bp paired end reads. RNA sequencing quality assurance was performed using FastQC (version 0.11.9), by looking for the presence of adapters and sequence quality. Genome alignment was mapped to the mouse genome (mm10-3.0.0) using 10x Genomics Cell Ranger pipeline (version 5.0.0). The resulting filtered feature-barcode matrices were used for downstream analysis. Data analysis and processing was performed using commercial code from Partek Flow package at https://www.partek.com/partek-flow/. GSEA was conducted using the R package Fast Gene Set Enrichment Analysis (fgsea) with KEGG, Reactome and Hallmark reference gene lists. The scRNA-seq data from P6 control retinas, extracted from our previous work and publicly available on NCBI’s Gene Expression Omnibus (accession no. GSE175895), were added to increase statistical power^29^.

### Statistical Analysis

All data are shown as mean± standard error of the mean (SEM) and were analyzed using the Student *t* test, 1-way ANOVA analysis of variance with the Sidak multiple comparison test, and 2-way analysis of variance with the Tukey multiple comparison test or the Sidak multiple comparison test. To construct the survival curves we have used the Kaplan–Meier method. All the statistical analyses were done using Prism 6 (GraphPad Software Inc). ns: nonsignificant, p>0.05, * p < 0.05, ** p < 0.01, *** p < 0.001.

## Results

### scRNA sequencing identifies signaling alterations in *Alk1* ECKO retinas

To identify gene expression changes in endothelial *Alk1* KO mice in an unbiased manner, we performed scRNA seq analysis of retinas from *Alk1^f/f^ Cdh5-CreERT2* and control mice at P6, following tamoxifen (TAM) injection at P4. ECs were isolated using CD31-coated magnetic beads and magnetic cell sorting (MACS) columns, employing a protocol we established previously^29^. A total of 21,277 cells from 6 pooled retinas per genotype were sequenced, with an average of 39,955 reads and 1,671 genes detected per cell. Low-quality cells and predicted doublets were removed prior to graph-based clustering and annotation based on gene expression patterns. Uniform manifold approximation and projection (UMAP) plots were generated for visualization, and cell populations were identified using established retinal cell markers. Approximately 10% of the retinal cells (2,352) were annotated as *Pecam1*+;*Cldn5*+ ECs (Figure 1A-1C), including distinct EC subtypes, such as venous (*Ptgis*^high^;*Nr2f2*^high^), proliferative (*Mki67*+;*Birc5*+), capillary (*Slc22a8*^high^;*Kcnj2*^high^), arterial (*Bmx*+;*Dkk2*+), and tip cell EC clusters (*Kcne3*+;*Angpt2*+)^29^. Notably, comparison between control and *Alk1* mutant ECs identified a specific EC cluster that was unique to *Alk1* mutants (Figure 1D-1F). This cluster potentially includes AVM-causing genes and was tentatively annotated as AVM EC. Venous cell-type signatures were most abundantly expressed in the AVM EC cluster, compared to arterial, capillary, and tip cell signatures (Figure 1F). Bioinformatic analysis of this cluster reveals upregulated PI3K/AKT/mTOR, VEGF, FOXO, Cell cycle signaling pathways, which we and others previously showed to be functionally involved in AVM formation, as well as fluid shear stress, hypoxia signaling, the complement cascade, and inflammatory signaling pathways (Figure 1G)^14, 16, 30-34^. Gene expression analysis of the AVM cluster compared to *Alk1* control showed decreased expression of *Alk1 (Acvrl1)*, *Smad4*, *Smad7*, *Plxnd1*, *Acvr2a*, *and Sema3a*, which are known to be downregulated with inhibition of ALK1 signaling (Figure 1H)^35-38^. In contrast, there was increased expression of *Hif1a* and *Slc2a1* (*Glut1*) (hypoxia signaling), *C1qa* (the complement cascade), *Piezo1*, *Klf4* and *Sqstm1* (*p62*) (fluid shear stress), and *Cd74* and *Lyz2* (inflammatory signaling pathways) (Figure 1 H). These data show that the loss of *Alk1* in ECs initiates distinct signaling pathways that could serve as therapeutic targets for treating AVMs.

**Figure 1.**
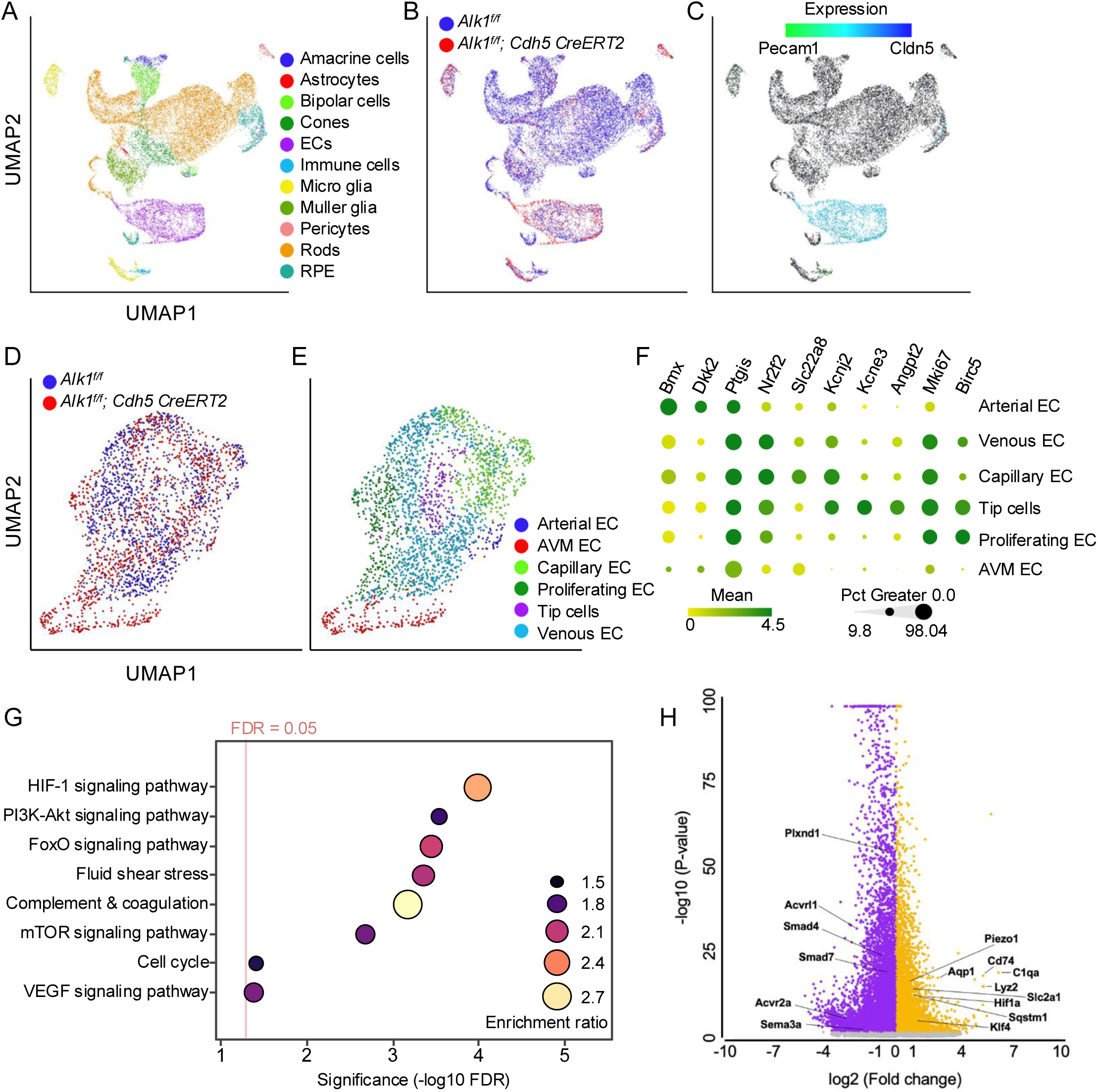
scRNA-seq analysis of *Alk1* ECKO. (A) UMAP plot of retinal cells from P6. (B) UMAP plot of retinal cells from P6 *Alk1* control and *Alk1* ECKO retinas. (C) UMAP plot of Pecam1 and Cldn5 from P6 *Alk1* control and *Alk1* ECKO retinal cells. (Pecam1: light green, Cldn5: blue) (D) UMAP plot of ECs from P6 *Alk1* control and *Alk1* ECKO retinas. (E) UMAP plot of EC sub-clusters from P6 *Alk1* control and *Alk1* ECKO retinas. (F) Dot plot of expression level and frequency among cell clusters of selected genes in P6 ECs. Color scale: green, high expression; yellow, low expression.(G) Gene set enrichment analysis of differentially expressed genes (DEGs) comparing the AVM cluster to *Alk1^f/f^* retinal ECs. (H) Volcano plot of DEGs between the AVM cluster in *Alk1* ECKO ECs and ECs from *Alk1* control at P6.

### PIEZO1 overexpression in ALK1 deficient endothelium in mice and HHT patients

We confirmed increased *Piezo1* mRNA expression levels in *Alk1* ECKO retinas compared to controls using qPCR (Figure 2A). Immunostaining with an antibody recognizing PIEZO1 revealed that *Alk1* ECKO upregulated endothelial PIEZO1 expression compared to TAM treated littermate controls (Figure 2B). PIEZO1 upregulation was observed throughout the *Alk1* deficient retina vasculature, including in AVMs (Figure 2B). The antibody specifically recognized PIEZO1, as immunostaining was abrogated in endothelial specific *Piezo1* knockout retinas (Supplementary Figure 1A). Moreover, *PIEZO1* mRNA and PIEZO1 protein expression increased in *ALK1* knockdown HUVECs compared to HUVECs transfected with a scrambled control siRNA (Figure 2C-2E), revealing that loss of endothelial ALK1 enhanced PIEZO1 expression in mouse and human ECs in a cell-autonomous manner.

**Figure 2.**
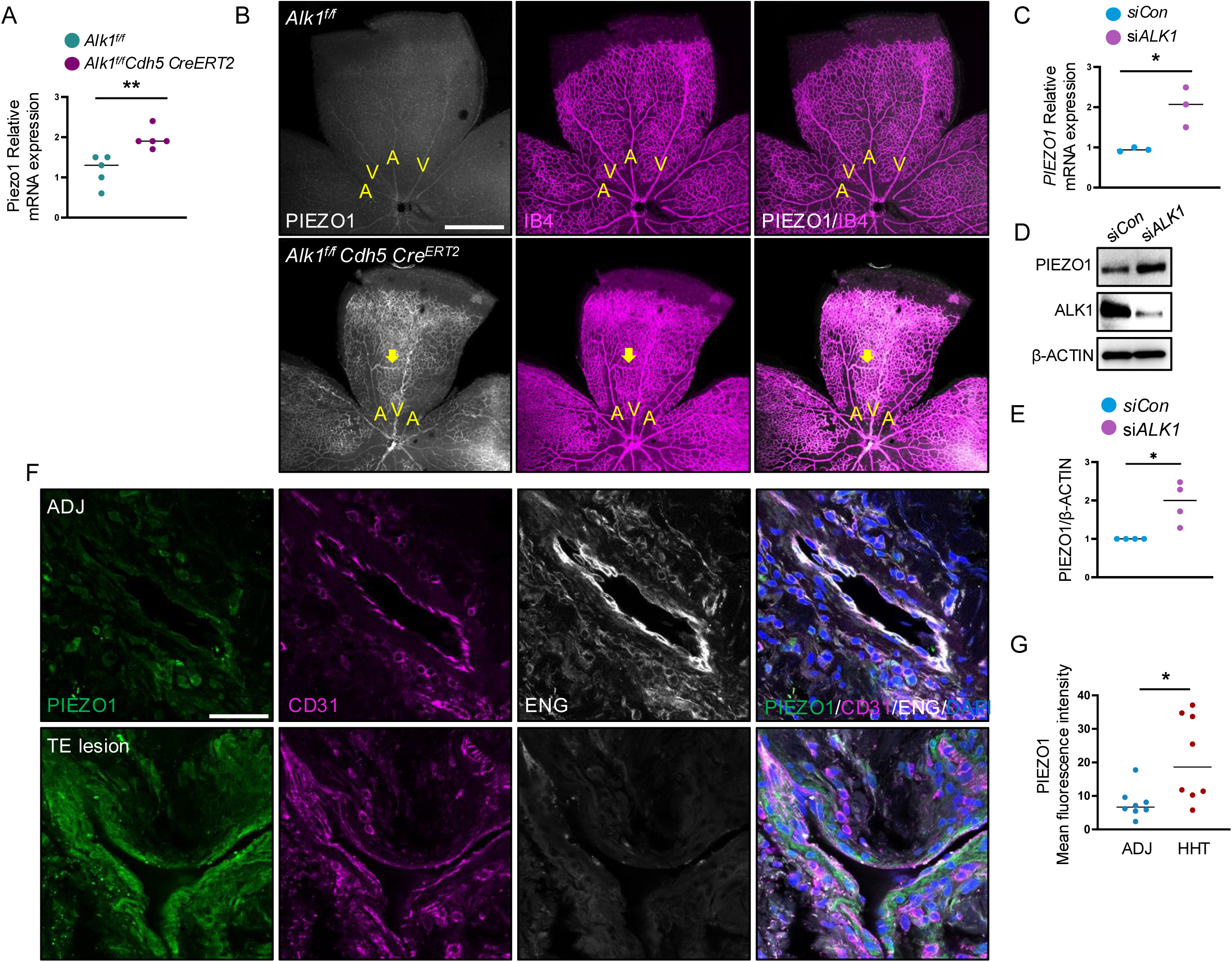
Upregulation of PIEZO1 in *Alk1* ECKO and human HHT2 patient samples. (A) PIEZO1 mRNA expression by qPCR in purified mouse retinas from P6 mice injected with 100 µg TAM at P4 (n=5). Error bars: SEM. **P-value < 0.01, Mann– Whitney *U* test. (B) IB4 (magenta) and PIEZO1 (white) staining of retinal flat mounts from postnatal day (P) 6 *Alk1^f/f^ Cdh5 Cre^ERT2^* and *Alk1^f/f^* pups. (C) *PIEZO1* mRNA expression by qPCR in HUVECs transfected with *Ctrl* or *ALK1* siRNA (n=3) (D) Western blot analysis HUVECs transfected with *Ctrl* or *ALK1* siRNA after 72 hr. (E) Quantifications of PIEZO1 levels normalized to β-ACTIN (n=4) (F) Immunostaining of healthy skin adjacent (ADJ) to telangiectasias and telangiectatic lesions (TE lesions) from patients with HHT type 2 for PIEZO1 (green), CD31 (magenta), and ENG (white). (G) Quantification of PIEZO1 staining (n=8; 4 patients, with 2 images selected per patient for analysis). Error bars: SEM. *P-value < 0.05, Mann–Whitney *U* test. Scale bars: 500 μm (B) and 50 μm (F).

To determine if PIEZO1 was also overexpressed in human HHT patients, we examined human dermal telangiectasia from patients with HHT type 2 and adjacent control normal skin biopsies. We observed increased PIEZO1 expression in dermal telangiectasias from HHT type 2 patients compared with normal skin samples (Figure 2F and 2G). PIEZO1 was overexpressed in CD31-positive ECs in the lesions, as well as by surrounding non-ECs. Expression of ENG in the telangiectasias was decreased (Figure 2F), confirming *ALK1* loss-of-function that reduces *Eng* expression as shown previously^8^.

### PIEZO1 function blocking in HHT2 mouse models

To test whether PIEZO1 inhibition could prevent AVM formation in *Alk1* ECKO mice, we intercrossed *Piezo1^f/f 39^*with *Alk1^f/f^ Cdh5 Cre^ERT2^*, a pan EC specific *Cre^ERT2^*. We also used *Alk1^f/f^ Mfsd2a Cre^ERT2^* mice^10^, in which *Alk1* deletion is driven by a venous and capillary EC specific *Cre^ERT2^*. We generated mice lacking one or both alleles of *Piezo1* in both *Alk1* mutant backgrounds and used TAM injection at P4 to delete both *Piezo1* and *Alk1* specifically in ECs (*Alk1^f/f^ Piezo1^f/f^ Cdh5 Cre^ERT2^*, *Alk1^f/f^ Piezo1^f/f^ Mfsd2a Cre^ERT2^*). Retinas were analyzed at P6 (Figure 3 A) and immunostaining of ALK1 and PIEZO1 confirmed the deletion of ALK1 and PIEZO1 in *Alk1^f/f^ Piezo1^f/f^ Cdh5 Cre^ERT2^* retinas (Supplementary Figure 1A). Heterozygous and homozygous deletion of *Piezo1* reduced AVM formation compared to *Alk1* ECKO with both Cre driver lines (Figure 3B and 3C, Supplementary Figure 1B and 1C). Homozygous deletion of *Piezo1* in *Alk1* ECKO backgrounds (*Alk1^f/f^ Piezo1^f/f^Cdh5 Cre^ERT2^* and *Alk1^f/f^ Piezo1^f/f^ Mfsd2a Cre^ERT2^*) retina showed even more reduced AVM vessel diameter and hypervascularization compared to *Alk1* ECKO and heterozygous deletion of *Piezo1* in *Alk1* ECKO retinas (Figure 3B and 3D and Supplementary Figure 1B and 1D).

**Figure 3.**
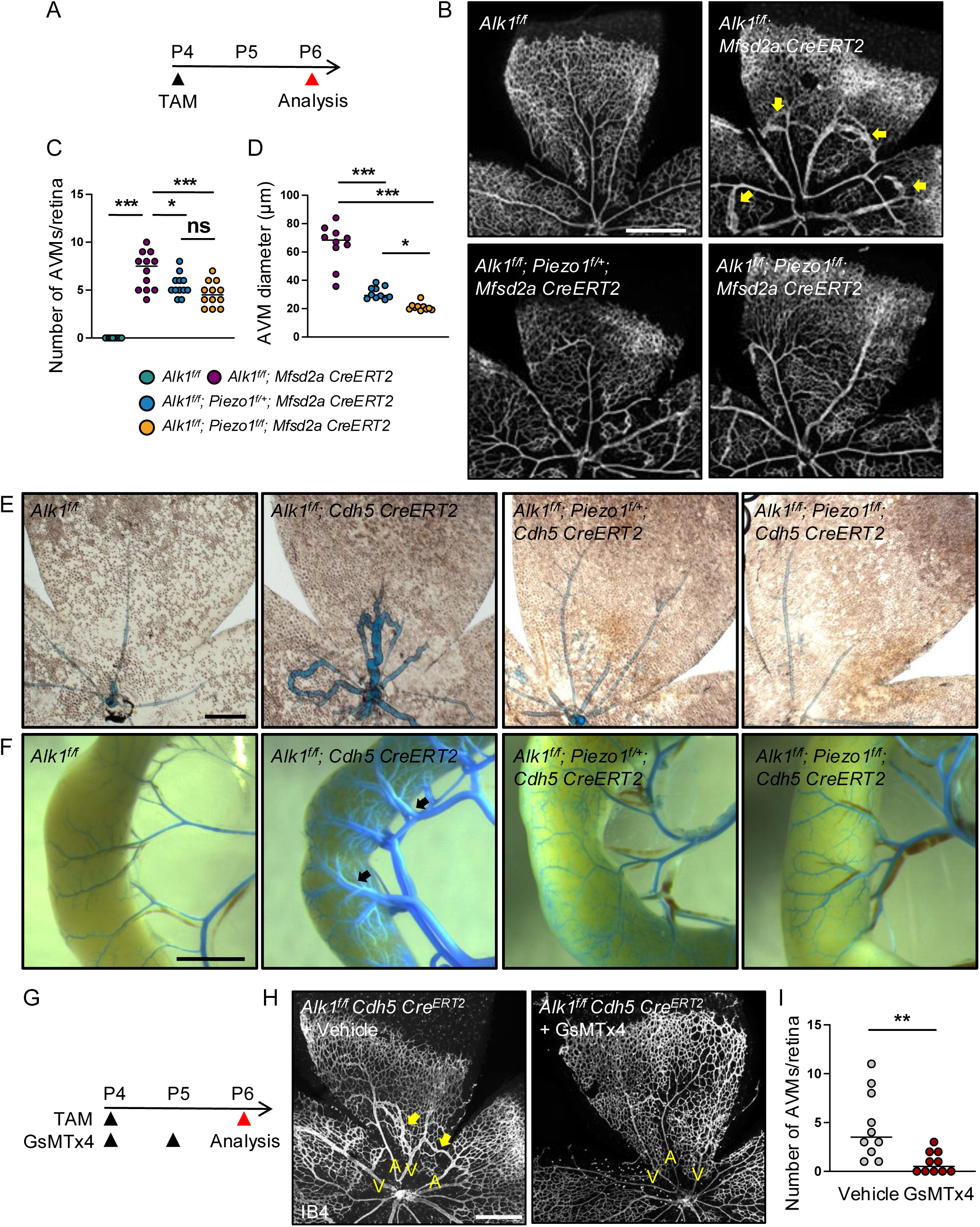
Inhibition of PIEZO1 prevents AVM formation in *Alk1* mutant mice. (A) Schematic representation of the experimental strategy for *Alk1* and *Piezo1* deletion in mice from P4 to P6. (B) IB4 staining of retinal flat mounts from *Alk1^f/f^* controls, *Alk1^f/f^ Mfsd2a Cre^ERT2^*, *Alk1^f/f^ Piezo1^f/+^ Mfsd2a Cre^ERT2^*, and *Alk1^f/f^ Piezo1^f/f^ Mfsd2a Cre^ERT2^* P6 mice. (C) Quantification of AVM number. n=10-12 mice per group. One-way ANOVA with Sidak’s multiple comparisons test. (D) Quantification of AVM vessel diameter (n=10-12). One-way ANOVA with Sidak’s multiple comparisons test. (E and F) Vascular labeling with latex dye (blue) in the retinas and gastrointestinal (GI) tract of *Alk1^f/f^* and *Alk1^f/f^ Cdh5 Cre^ERT2^*, *Alk1^f/f^ Piezo1^f/+^ Cdh5 Cre^ERT2^*, and *Alk1^f/f^ Piezo1^f/f^ Cdh5 Cre^ERT2^* P6 pups. Black arrows indicate AVMs. (G) Experimental strategy to assess the effects of PIEZO1 inhibitor in *Alk1* deleted retinas. Arrowheads indicate the time course of TAM (100 μg) and GsMTx4 (1 mg/kg) or vehicle administration. (H) IB4 staining of P6 retinal flat mounts from *Alk1^f/f^ Cdh5 Cre^ERT2^* mice injected with GsMTx4 or vehicle at P4 and P5. ‘A’ indicates arteries, ‘V’ indicates veins, and arrows denote AVMs. (I) Quantification of AVM count. Each dot represents one retina (n=10). Mann–Whitney U test. Error bars: SEM. *P-value < 0.05, **P-value < 0.01, *** P-value < 0.001, ns: nonsignificant, Scale bars: 500 μm (B and H), 200 μm (E), 10 mm (F)

In addition to IsoB4 staining to visualize direct artery-vein connections, we used intracardiac latex dye injection in deeply anesthetized mice. The latex dye does not cross the capillary beds and was retained within the arterial branches in *Alk1^f/f^* retina and gastrointestinal (GI) tract (Figure 3E and 3F). In the *Alk1* ECKO, the latex penetrated both the venous and the arterial branches via AVMs in the retina and GI tract (Figure 3E and 3F). The frequency of AVMs and latex perfused veins in double *Alk1 Piezo1* ECKO mice was reduced when compared to the *Alk1* ECKO pups (Figure 3E and 3F). We further tested whether double *Alk1 Piezo1* ECKO could enhance the survival rate of *Alk1* ECKO mutants. It is known that *Piezo1* KO mice showed lethality due to complications of development and homeostasis^23, 24^. However, heterozygous mutation of *Piezo1* in *Alk1^f/f^ Mfsd2a Cre^ERT2^* mutants prolonged survival by 2-3 days and homozygous mutations of *Piezo1* in *Alk1^f/f^ Mfsd2a Cre^ERT2^* mutants prolonged survival by 5-6 days longer survival rates compared to *Alk1^f/f^ Mfsd2a Cre^ERT2^* (Supplementary Figure 1E).

To test pharmacological interventions based on PIEZO1 inhibition, we used a known PIEZO1 antagonist GsMTx4, which is a gating modifier peptide from spider venom that selectively inhibits cation-permeable mechano-sensitive channels such as PIEZO1 and is widely used for PIEZO1 function inhibition^21, 23, 40-43^. *Alk1* deletion was induced by TAM injection at P4, inhibitors were given intraperitoneally at 1 mg/kg P4 and P5, and mice were analyzed at P6 (Figure 3G). GsMTx4 decreased AVM formation in *Alk1^f/f^ Cdh5 Cre^ERT2^* mice (Figure 3H and 3I). Collectively, these data indicate that PIEZO1 overexpression contributes to AVM formation in *Alk1* mutants, and that its inhibition can reduce AVMs in mouse mutants.

### PIEZO1 lies upstream of the VEGFR2 and ERK5 signaling pathways

As PIEZO1 is implicated in LSS mechanotransduction^19, 20^, and AVM formation in *Alk1* mutants is regulated by flow^10, 13, 44^, we tested if ALK1-PIEZO1 signaling effects were shear stress dependent. HUVECs were transfected with either scrambled control siRNA, *ALK1* siRNA, *PIEZO1* siRNA or both *ALK1 and PIEZO1* siRNAs and cultured in static conditions or under shear stress. Protein extracts from these cells were analyzed by western blot with antibodies against PIEZO1, VEGFR2 and downstream effectors. PIEZO1 protein levels were significantly increased by ALK1 knockdown in both static and flow conditions (Figure 4A -4D). Next, we assessed VEGFR2 expression levels and the phosphorylation of VEGFR2 at Y1175, which is enhanced by shear stress and triggers downstream signaling^45, 46^. Total VEGFR2 expression was significantly increased in ALK1-depleted ECs in static conditions and increased even more under flow conditions (Figure 4A and 4B). Notably, VEGFR2 Y1175 phosphorylation was enhanced in *ALK1*-depleted ECs under flow but was normalized in cells treated with both *ALK1* and *PIEZO1* siRNAs (Figure 4A and 4B).

**Figure 4.**
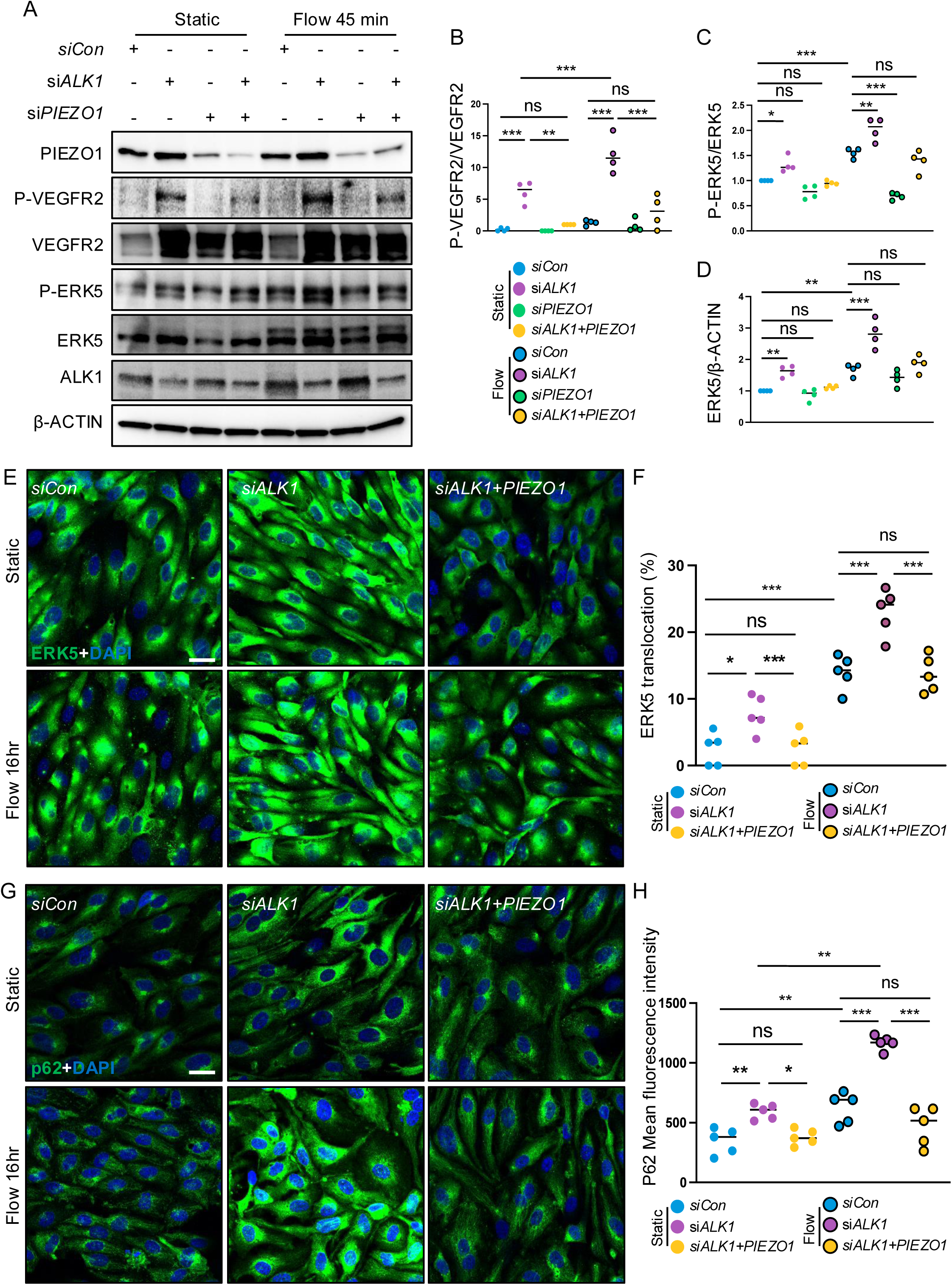
PIEZO1 functions upstream of ALK1-mediated VEGFR2-ERK5 signaling pathway. (A) Western blot analysis of HUVECs transfected with control, *ALK1*, *PIEZO1*, and *ALK1*+*PIEZO1* siRNAs, followed by 45 min of flow exposure. (B-D) Quantification of P-VEGFR2/VEGFR2, P-ERK5/ERK5 and ERK5/β-ACTIN. One-way ANOVA with Sidak’s multiple comparisons test. (E) ERK5 staining for *siCon*, *siALK1* and *siALK1*+*siPIEZO1* transfected HUVECs under static conditions and after 16 hr of flow exposure. ERK5 (green) DAPI (blue) (F) Quantification of nuclear ERK5 translocation for *siCon*, *siALK1* and *siALK1*+*siPIEZO1* transfected HUVECs under static conditions and following 16 hr of flow exposure. *P<0.05, ***P<0.001, ns: nonsignificant, One-way ANOVA with Sidak’s multiple comparisons test. (G) p62 immunostaining in *siCon*, *siALK1* and *siALK1*+*siPIEZO1* transfected HUVECs under static conditions and following 16 hr of flow exposure. (H) Quantification of p62 expression in *siCon*, *siALK1* and *siALK1*+*siPIEZO1* transfected HUVECs under static conditions and after 16 hr of flow exposure. One-way ANOVA with Sidak’s multiple comparisons test. *P<0.05, **P<0.01, ***P<0.001, ns: nonsignificant, Scale bars: 25 μm (E and G)

One way in which PIEZO1 regulates mechanotransduction is through activation of phospho-ERK5 and ERK5^19, 20^. Interestingly, total ERK5 expression was upregulated in *ALK1*-depleted ECs, with an even greater increase observed under flow conditions (Figure 4A and 4D). Laminar flow-exposed cells exhibit a phosphorylation-dependent mobility shift ("gel shift") of the total ERK5 protein, indicating ERK5 activation^47^, which corresponds to autophosphorylation in the C-terminal region^48, 49^ (Figure 4A). Phosphorylation at Thr218 and Tyr220 of ERK5 within the TEY motif is selectively mediated by its sole upstream kinase, MEK5^50^. To detect the MEK5 specific-phosphorylation, we used an antibody specific to phospho-ERK5 (Thr218/Tyr220). Notably, ERK5 phosphorylation was markedly elevated in *ALK1*-depleted ECs, attenuated in PIEZO1-knockdown ECs, and normalized in *ALK1* and *PIEZO1* double-knockdown ECs under flow conditions compared to static conditions (Figures 4A and 4C). Phosphorylated ERK5 is known to translocate to the nucleus, where it activates KLF4^47,51^. Immunostaining of ERK5 demonstrated increased nuclear translocation in *ALK1*-depleted HUVECs, while nuclear translocation was diminished in *ALK1* and *PIEZO1* double-depleted HUVECs under static conditions. Under flow conditions, nuclear translocation of ERK5 was further elevated in *ALK1*-depleted HUVECs and normalized in *ALK1* and *PIEZO1* double-knockdown HUVECs (Figure 4E and 4F). We also investigated p62, a downstream target of ERK5 that was upregulated in the AVM cluster identified from scRNA-seq analysis. Immunostaining revealed increased p62 expression in *ALK1*-depleted HUVECs, but reduced in *ALK1* and *PIEZO1* double-knockdown HUVECs under static conditions (Figure 4G and 4H). Under flow conditions, p62 expression was further elevated in *ALK1*-depleted HUVECs, exhibiting punctate speckling and was normalized in *ALK1* and *PIEZO1* double-knockdown HUVECs (Figure 4G and 4H). These findings indicate that PIEZO1 functions upstream of VEGFR2 and ERK5 within the flow induced ALK1 signaling pathway.

### KLF4 overexpression in ALK1 deleted ECs and human HHT2

As KLF4 was upregulated in the *Alk1* mutant scRNA-seq AVM cluster (Figure 1H), and lies downstream of the PIEZO1-ERK5-p62 signaling pathway^19, 20^, we tested *Klf4* mRNA and KLF4 protein expression in *Alk1^f/f^ Cdh5 Cre^ERT2^* retinas. We confirmed increased *Klf4* mRNA expression levels in *Alk1* ECKO retinas compared to controls using qPCR (Supplementary Figure 2A). Immunostaining with a knockout validated antibody showed faint KLF4 expression in arterioles, venules, and capillaries proximal to the optic nerve in *Alk1^f/f^* retinas (Supplementary Figure 2B). However, in *Alk1^f/f^ Cdh5 Cre^ERT2^* retinas, KLF4 expression was elevated specifically in AVM vessels (Supplementary Figure 2B). Additionally, genetic deletion of *Piezo1* in *Alk1^f/f^ Mfsd2a Cre^ERT2^* mice demonstrated reduced and normalized KLF4 expression in *Alk1^f/f^ Piezo1^f/f^ Mfsd2a Cre^ERT2^* retinas (Figure 5A). To test KLF4 expression was affected by pharmacological inhibition of PIEZO1 *in vivo*, we inhibited PIEZO1 using GsMTx4 in *Alk1^f/f^ Cdh5 Cre^ERT2^* mice. GsMTx4-injected *Alk1* ECKO retinas exhibited reduced KLF4 expression compared to untreated *Alk1* ECKO retinas (Supplementary Figure 2C).

**Figure 5.**
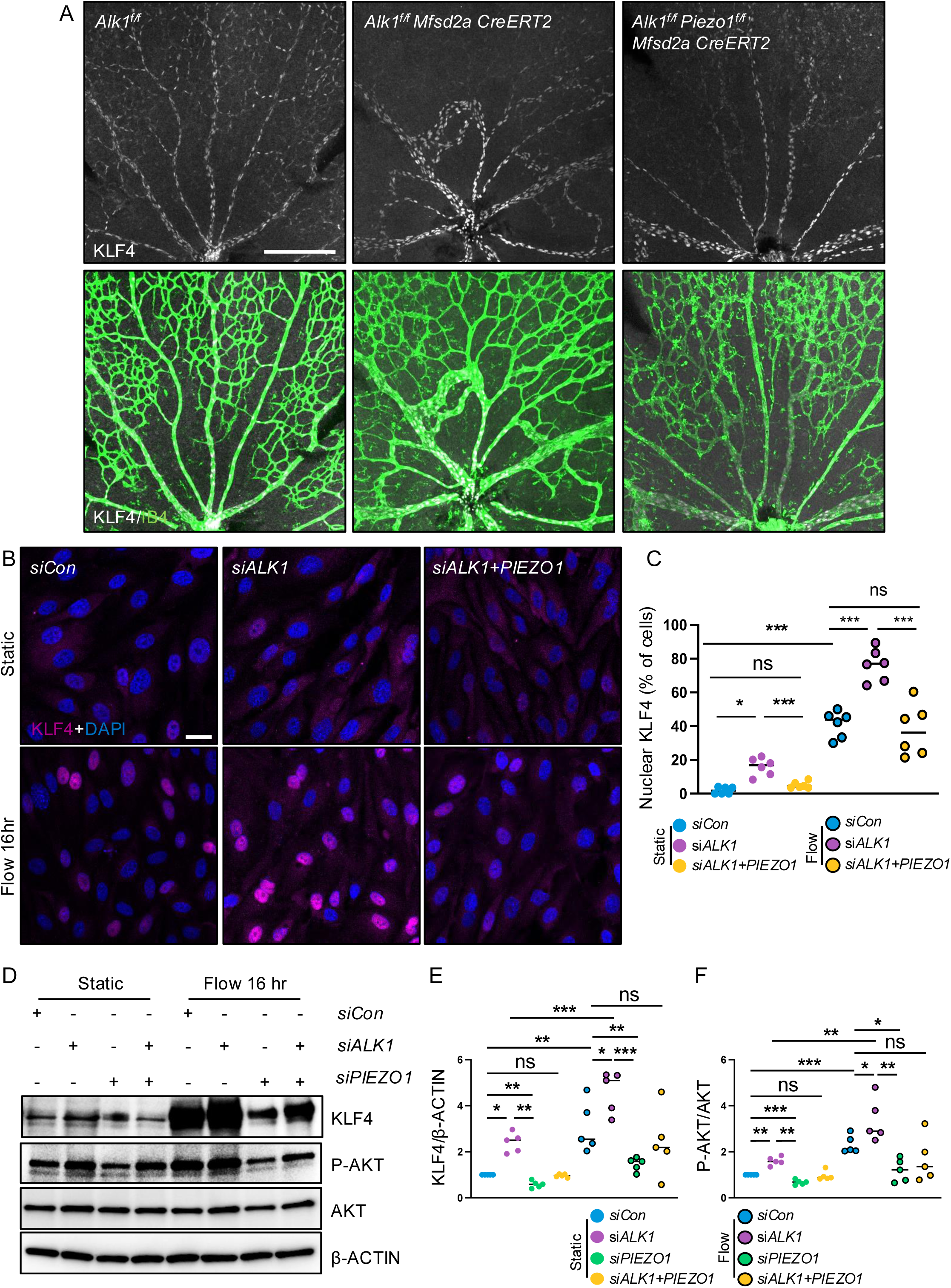
PIEZO1 inhibition prevents KLF4 upregulation in *Alk1* mutant retinas. (A) KLF4 (white) and IB4 (green) staining of retinal flat mounts from P6 *Alk1^f/f^*, *Alk1^f/f^ Mfsd2a Cre^ERT2^* and *Alk1^f/f^ Piezo1^f/f^ Mfsd2a Cre^ERT2^* pups. (B) KLF4 immunostaining in *siCon*, *siALK1* and *siALK1*+*siPIEZO1* transfected HUVECs under static conditions and after 16 hr of flow exposure. KLF4 (magenta), DAPI (blue) (C) Quantification of nuclear KLF4 in HUVECs transfected with siCon, si*ALK1*, or si*ALK1*+si*PIEZO1* under static conditions and following 16 hr of flow exposure. One-way ANOVA with Sidak’s multiple comparisons test. (D) Western blot analysis of HUVECs transfected with control, *ALK1*, *PIEZO1*, and *ALK1*+*PIEZO1* siRNAs, followed by 16 hr of flow exposure. (E and F) Quantification of KLF4/β-ACTIN and P-AKT/AKT. One-way ANOVA with Sidak’s multiple comparisons test. *P<0.05, **P<0.01, ***P<0.001, ns: nonsignificant, Scale bars: 250 μm (A) and 25 μm (B)

Next, to determine whether KLF4 was involved in ALK1 signaling, we assessed nuclear KLF4 expression in control, *ALK1*-knockdown, and *ALK1* and *PIEZO1* double-knockdown ECs, both under static conditions and following exposure to flow for 16 hr. Nuclear KLF4 expression was increased in *ALK1*-depleted ECs but reduced in *ALK1* and *PIEZO1* double-knockdown ECs under static conditions (Figure 5 B and 5C). Under flow conditions, nuclear KLF4 expression was enhanced in control siRNA-treated ECs compared to static conditions (Figure 5 B and 5C). KLF4 expression was significantly elevated in *ALK1*-depleted ECs under flow conditions compared to flow-induced control siRNA-treated ECs, and reduced in both *PIEZO1* knockdown and *ALK1* and *PIEZO1* double-knockdown ECs under flow conditions (Figure 5B and 5C). Western blot analysis of HUVECs cultured as described above confirmed the upregulation of KLF4 protein levels in *ALK1*-depleted cells, the reduction of KLF4 protein levels in *PIEZO1*-depleted cells and the normalization of KLF4 protein levels in *ALK1-PIEZO1* double-knockdown cells under both static and flow conditions (Figure 5D and 5E). Interestingly, AKT activation (S473) was elevated in *ALK1*-depleted ECs but reduced in *PIEZO1*-depleted ECs. Additionally, *ALK1* and *PIEZO1* double-knockdown ECs exhibited decreased AKT activation compared to *ALK1*-depleted ECs. Under flow conditions, AKT phosphorylation was enhanced in control siRNA-treated ECs compared to static conditions (Figure 5D and 5F). Notably, AKT phosphorylation was further increased in flow-mediated *ALK1*-depleted ECs but was reduced in *ALK1* and *PIEZO1* double-depleted cells (Figure 5C and 5E). Lastly, we examined KLF4 and PIEZO1 expression in human dermal telangiectasias from patients with HHT type 2, and observed increased nuclear KLF4 expression as well as PIEZO1 overexpression in CD31-positive ECs and non ECs in the lesions compared to adjacent control normal skin biopsies (Figure 6A and 6B). Collectively, these findings implicate upregulation of the PIEZO1-ERK5-p62-KLF4 pathway in ALK1 signaling in mice and humans.

**Figure 6.**
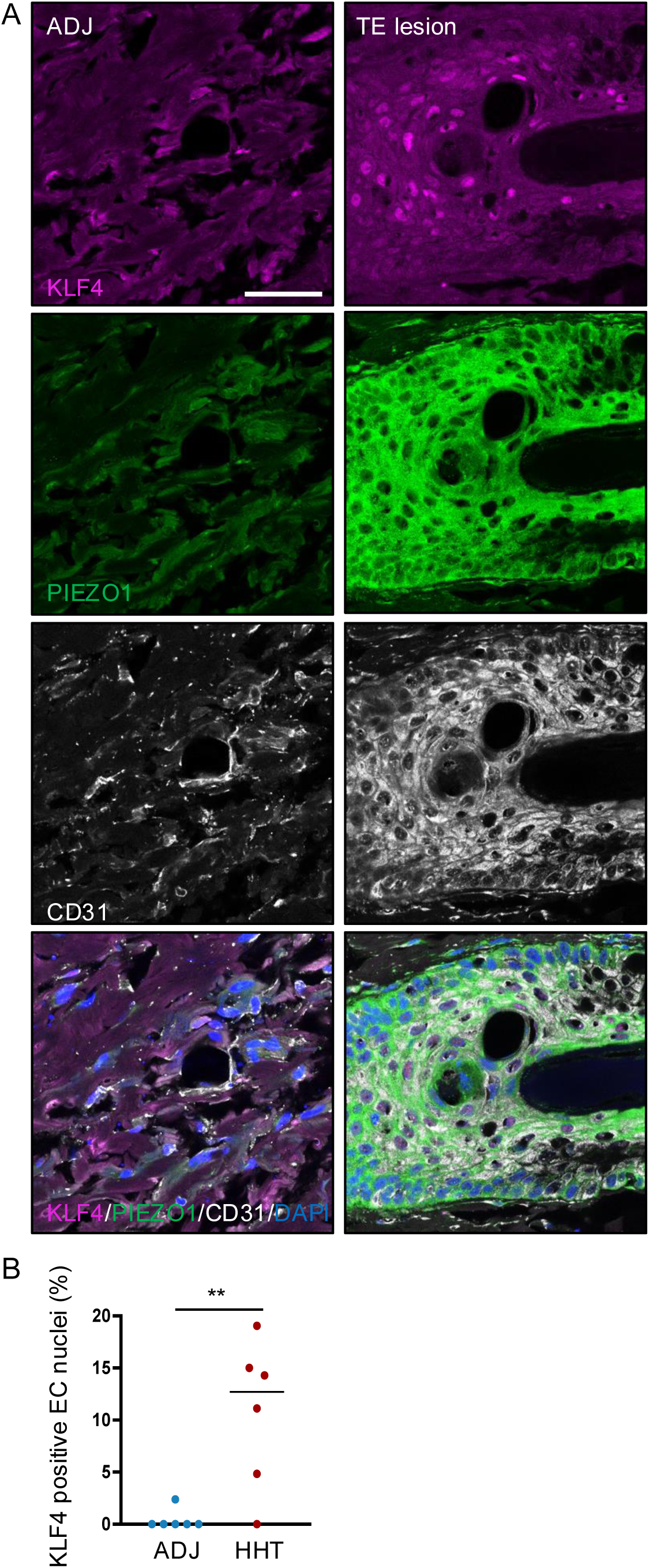
Upregulation of KLF4 and PIEZO1 in human HHT2 patient samples. (A) Immunostaining of healthy skin adjacent (ADJ) to telangiectasias and telangiectatic lesions (TE lesions) from patients with HHT type 2 for KLF4 (magenta), PIEZO1 (green), and CD31 (white). (B) Quantification of KLF4 staining (n=6; 3 patients, with 2 images selected per patient for analysis). Error bars: SEM. *P-value < 0.05, Mann–Whitney *U* test. Scale bar: 50 μm (A)

### PIEZO1 inhibition rescues hypoxia in *Alk1* ECKO mice

Given that hypoxia and HIF1 signaling were elevated in the *Alk1* mutant scRNA AVM cluster, we aimed to determine whether PIEZO1 inhibition could mitigate hypoxia in *Alk1* mutant mice. First, we assessed hypoxia using pimonidazole *in vivo*. Pimonidazole is converted by hypoxia-activated nitroreductases into a reactive intermediate that covalently binds to cellular components, forming adducts in hypoxic regions^52^. Mice were injected with 60 mg/kg of pimonidazole for 90 minutes, after which pimonidazole levels were detected using anti-pimonidazole adduct antibody. *Alk1^f/f^* mice stained positive for pimonidazole in the non-vascularized, distal regions of the retinas, while the vascularized and oxygenated retina stained negative as expected^53^ (Figure 7A). By contrast, in *Alk1^f/f^ Cdh5 Cre^ERT2^* mice pimonidazole was detected throughout the entire retina (Figure 7A). *Alk1 Piezo1* double mutant ECKO retinas exhibited pimonidazole staining that was similar to *Alk1^f/f^* mice and reduced compared to *Alk1^f/f^ Cdh5 Cre^ERT2^* (Figure 7A). To directly evaluate HIF1A expression levels, we immunolabeled HIF1A in the retinas. HIF1A expression was detected in non-vascularized, distal regions of the retinas in *Alk1*^fl/fl^ mice, upregulated in the vasculature of *Alk1^f/f^ Mfsd2a Cre^ERT2^* retinas, including in AVM ECs, and normalized in *Alk1^f/f^ Piezo1^f/f^ Mfsd2a Cre^ERT2^* retinas to levels that were comparable to *Alk1*^fl/fl^ retinas (Figure 7B), thus mirroring results obtained with Pimonidazole.

**Figure 7.**
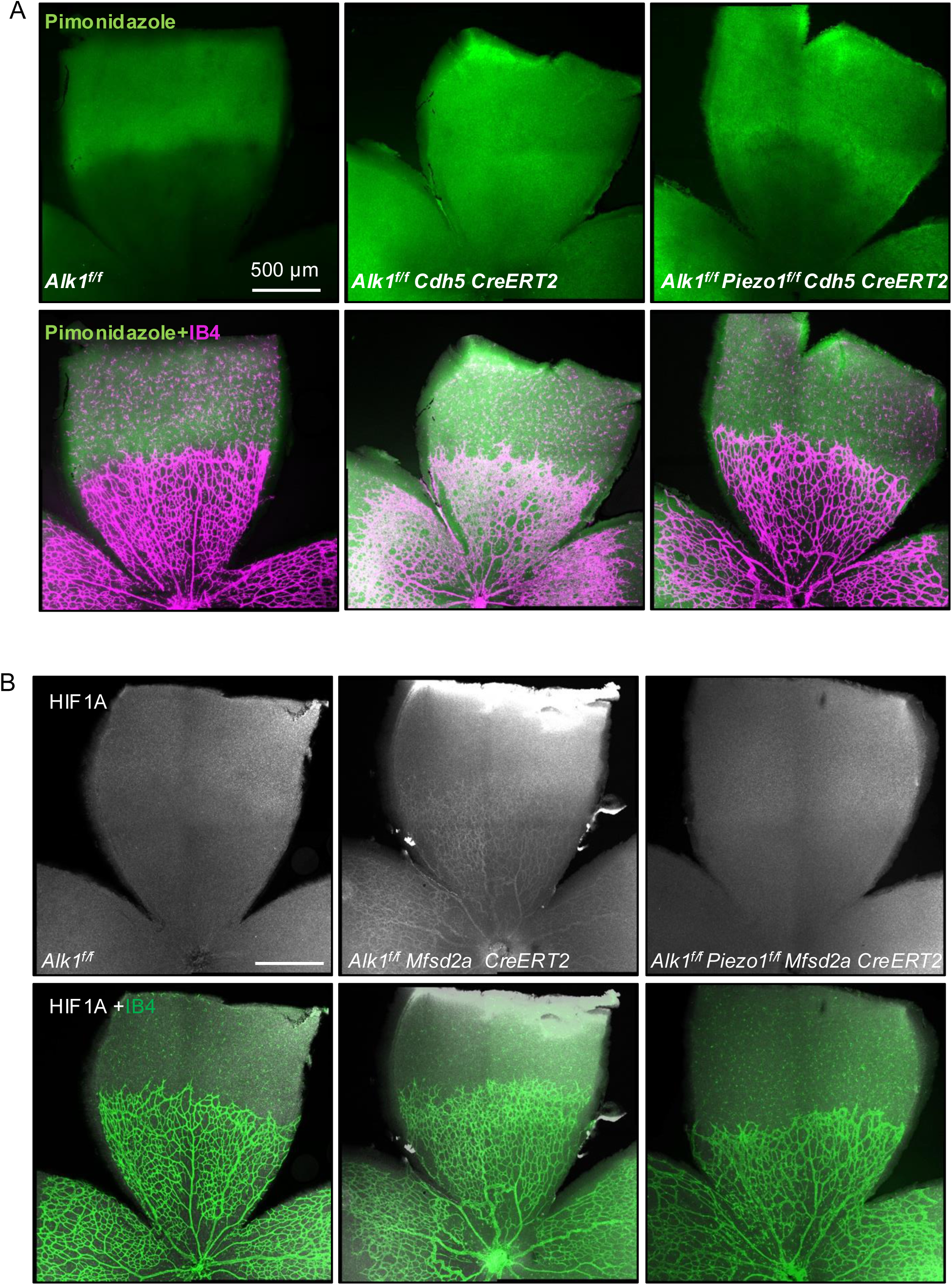
PIEZO1 inhibition blocks hypoxia in *Alk1* mutant retinas. (A) Area of retinal hypoxia 90 min after pimonidazole injection in *Alk1^f/f^*, *Alk1^f/f^* Cdh5 *Cre^ERT2^* and *Alk1^f/f^ Piezo1^f/f^ Cdh5 Cre^ERT2^* P6 pups. Hypoxyprobe (green), IB4 (magenta) (B) Immunostaining of HIF1A (white) and IB4 (green) in retinal flat mounts from P6 *Alk1^f/f^*, *Alk1^f/f^ Mfsd2a Cre^ERT2^* and *Alk1^f/f^ Piezo1^f/f^ Mfsd2a Cre^ERT2^* mice. Scale bars: 500 μm (A and B),

### PIEZO1 inhibition rescues EC proliferation in *Alk1* ECKO mice

We and others had demonstrated that increased EC proliferation contributed to AVM foramation^14, 31, 33, 34^, and scRNA-seq analysis confirmed an upregulation of cell cycle genes in the AVM cluster. To determine whether *Piezo1* deletion could prevent hyperproliferation in *Alk1*-mutant ECs, we administered EdU 4 hr before analysis to label proliferating cells. Compared to *Alk1^f/f^* retinas, the vascular plexus of *Alk1^f/f^ Mfsd2a Cre^ERT2^* retinas exhibited an increased number of ERG+ EdU+ double positive cells, whereas *Alk1^f/f^ Piezo1^f/f^ Mfsd2a Cre^ERT2^* showed a reduction in ERG+ EdU+ double-positive cells compared to *Alk1^f/f^ Mfsd2a Cre^ERT2^* retinas (Figure 8A and 8B). Furthermore, we tested whether PIEZO1 inhibition could mitigate the increased cell proliferation seen in ALK1-deficient HUVECs using EdU incorporation assay. ECs with dual *ALK1* and *PIEZO1* depletion exhibited reduced proliferation compared to ECs with *ALK1* depletion alone (Figure 8 C and 8D). These findings suggest that PIEZO1 plays a role in cell proliferation within the context of ALK1 signaling.

**Figure 8.**
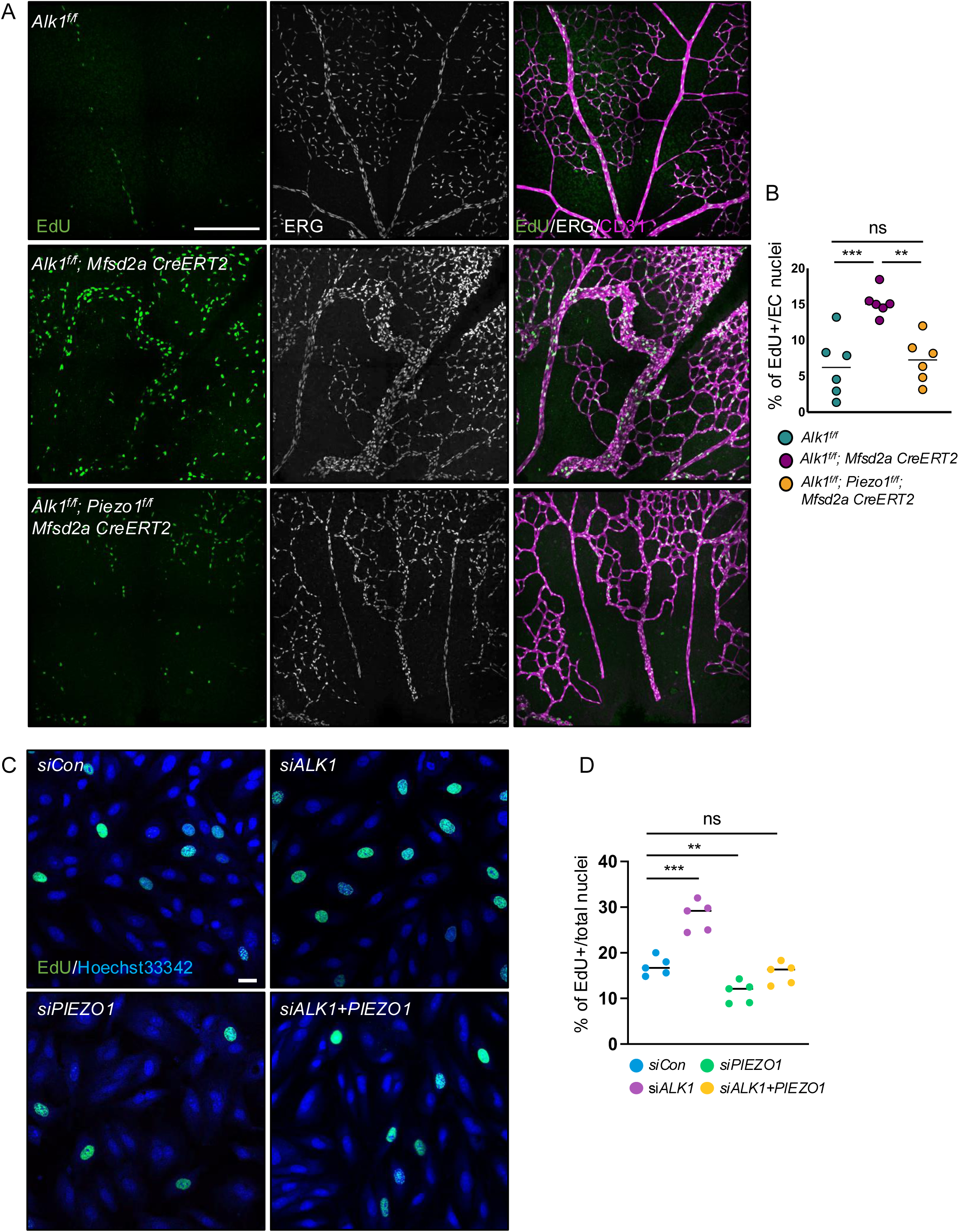
PIEZO1 inhibition blocks hyperproliferation in ALK1-depleted ECs. (A) Labeling for EdU (green), ERG (white), and CD31 (magenta) in the vascular plexus of retinas from P6 *Alk1^f/f^* and *Alk1^f/f^ Mfsd2a Cre^ERT2^* and *Alk1^f/f^ Piezo1^f/f^ Mfsd2a Cre^ERT2^* mice. (B) Quantification of the number of ERG and EdU double positive nuclei per vascular area. Mann–Whitney U test. (C) Labeling for EdU (green) and Hoechst33342 (blue) in HUVECs transfected with control, *ALK1*, *PIEZO1*, and *ALK1*+*PIEZO1* siRNAs. (D) Quantification of the number of EdU and Hoechst33342 double positive nuclei (n = 5). Two-way ANOVA with Tukey’s multiple comparisons test. Error bars: SEM. **P-value < 0.01, ***P<0.001, ns: nonsignificant, Scale bars: 500 μm (A), 25 μm (C)

## Discussion

Our scRNA-seq gene set enrichment analysis revealed that the complement cascade, hypoxia, inflammatory signaling, and high fluid shear stress were among the most upregulated pathways in *Alk1* KO ECs, along with previously identified signaling alterations, such as cell cycle, PI3K/AKT, mTOR, FOXO, and VEGF signaling. Of note, scRNA-seq data from spontaneous human brain AVMs^30^ show similar gene set enrichment, suggesting that upregulation of these pathways is common between spontaneous AVMs and in HHT, thereby broadening the impact of our study to an even wider patient population.

We identified a novel regulatory arm of ALK1 signaling, wherein the loss of ALK1 enhances PIEZO1 expression and PIEZO1-mediated VEGFR2-PI3K-AKT signaling, leading to increased KLF4 signaling. In the absence of ALK1, elevated flow-induced PIEZO1 activation triggers both VEGFR2 signaling and the p62-ERK5 mitophagy pathway, resulting in increased KLF4 expression and nuclear localization. Notably, both pharmacological and genetic inhibition of PIEZO1 effectively prevented vascular malformations in *Alk1*-deficient mice. These findings highlight the critical role of PIEZO1 as a mechanosensor within the ALK1 pathway and suggest a potential therapeutic target for managing vascular abnormalities in HHT.

The present study reveals a novel signaling pathway involving flow mediated PIEZO1-ERK5-p62-KLF4 overactivation in the context of HHT. We demonstrated increased PIEZO1 and KLF4 expression in human dermal telangiectasias from HHT type 2 patients, as well as in *Alk1* endothelial knockout mouse models. Mechanistically, ALK1 deletion led to elevated PIEZO1 and p62 expression, which in turn activated ERK5 and promoted nuclear KLF4 expression. Importantly, both pharmacological inhibition and genetic deletion of PIEZO1 mitigated AVM formation and reduced hypoxia-cell proliferation associated signaling.

The mitochondrial pathway is activated by the PIEZO1, which increased oxidative phosphorylation and reactive oxygen species generation^54^. This damages mitochondria and initiates mitophagy by stabilization of PINK1, in turn resulting in assembly of a p62-dependent scaffolding complex that amplifies MEKK–MEK5–ERK5 signaling and enhanced flow-induced KLF2 expression. p62 or PINK1 KO mice showed reduced expression of *Klf2* and KLF2-dependent genes and inhibited FSS-mediated vascular remodeling^55^. Strikingly, beyond PIEZO1, several key components of the mitophagy pathway were significantly upregulated in the *Alk1* ECKO AVM EC cluster. These include *Smdt1*, encoding EMRE, a subunit of the mitochondrial calcium uniporter essential for calcium entry into mitochondria, and *Sqstm1*, encoding p62, a mitophagy adapter protein critical for ERK5 activation through the mitochondrial pathway, as well as *Klf4* itself. The structural and functional abnormalities associated with AVMs disrupt normal blood flow, resulting in insufficient delivery of oxygen- and nutrient-rich blood. These vascular defects induce a pathological microenvironment characterized by hypoxia, inflammation, and nutrient deprivation. In response, flow-mediated PIEZO1 activation triggers the MEK5-ERK5-p62-KLF2/4mitophagy pathway in ECs as a compensatory mechanism to maintain vascular homeostasis^19, 56-58^. We found that elevated PIEZO1 in *Alk1* mutants contributes to AVM formation by overactivating the mitophagy pathway and KLF4 expression, resulting in aberrant endothelial gene expression that drives abnormal cellular behavior. Notably, recent work demonstrated that *Smad4* knockout or knockdown (KO/KD) sensitizes ECs to FSS, allowing for alignment and KLF4 induction at much lower FSS levels than in control cells, directly supporting the notion that HHT involves amplified FSS signaling^32^. Thus, our findings establish that dysregulated shear stress sensing is a central mechanism in AVM formation. However, further investigation is required to elucidate the detailed role of the mitophagy pathway in the pathogenesis of HHT.

In two different genetic backgrounds, *Piezo1* endothelial knockout (KO) mice exhibited embryonic lethality during mid-gestation, displaying growth retardation and impaired vascular remodeling in the yolk sac^23^. Another model of *Piezo1* endothelial conditional KO mice survived until approximately two weeks after birth but subsequently died, presenting with significant vascular defects^24^. These findings underscore a critical role for endothelial PIEZO1 in vascular development and postnatal survival. In TAM-inducible *Alk1* pan-endothelial KO mice, TAM administration at P4 resulted in mortality within 48 hr, whereas venous endothelial-specific *Alk1* KO mice survived 4–6 days post-TAM induction at the same age. Notably, double KO of *Alk1* and *Piezo1* in venous endothelial cells led to extended survival compared to *Alk1* KO mice, with mice living for 10–12 days post-TAM injection; however, they eventually succumbed due to the compounded effects of *Piezo1* and *Alk1* deficiencies. These results highlight the complex interplay between PIEZO1 and ALK1 in ECs, suggesting that both are essential for proper vascular function and survival beyond early development.

Our data show that increased hypoxia signaling pathway in absence of ALK1 was normalized by PIEZO1 deletion in retinal AVMs in *Alk1* ECKO mice. Hypoxia is a hallmark of AVMs and has been associated with the upregulation of HIF1A and GLUT1, as observed in human AVM samples. Consistently, our scRNA-seq analysis revealed increased *Hif1a* and *Glut1* within the AVM cluster in *Alk1* ECKO mouse retinas. Previous studies have shown that HIF1A is expressed in cerebral AVM, accompanied by elevated levels of VEGF and VEGFRs^59, 60^. Interestingly, while extracranial AVMs lack GLUT1 expression, GLUT1/CD31 colocalization has been observed in ECs within cerebral AVM and CCM samples^61^. Hypoxia is known to induce PIEZO1 expression in ECs, as demonstrated in pulmonary arterial ECs from patients with idiopathic PAH, known to be hypoxic, showed increased PIEZO1 expression^62^. Furthermore, hypoxia has been proposed as a potential non-genetic second hit contributing to AVM development in HHT ^63, 64^. For the first time, our data show that endothelial HIF1A expression is elevated in the retinas of *Alk1* ECKO mice and is normalized in *Alk1* and *Piezo1* double ECKO mouse retinas, highlighting the critical interplay between ALK1, PIEZO1, and hypoxia signaling in AVM pathophysiology.

Increased cell proliferation, indicated by Ki67 staining, vessel enlargement, and higher expression of extracellular matrix molecules have been observed in cutaneous telangiectasia biopsies from human patients with HHT1 and HHT2 compared to controls^16,31^. Recent studies, along with our own data, demonstrate hyperproliferation in the retinal ECs of *Alk1* and *Smad4* ECKO mouse models within AVM regions. Importantly, palbociclib, a cell cycle inhibitor, effectively prevented AVM formation in BMP9/10-blocking antibody-treated mice and *Alk1* ECKO mice^14, 31, 33, 34^. In our study, ALK1 and PIEZO1 double-depleted ECs exhibited reduced cell proliferation compared to ALK1-depleted ECs, both *in vitro* and *in vivo*. Here, we extend these findings by demonstrating that PIEZO1 is essential for proper vessel development, regulating cell cycle progression and hypoxia response upstream of the ALK1-mediated VEGFR2-PI3K signaling pathway in ECs.

To date, current therapeutic options for HHT primarily aim to alleviate disease symptoms, such as epistaxis. Both preclinical and clinical studies exploring anti–VEGF-A, anti-ANG2 molecules, and PI3K inhibitors are emerging as potential treatment^65-68^. These therapies aim to target the overactivated proangiogenic pathways in HHT, with the goal of restoring AVMs to normal vasculature. A recent study demonstrated that pomalidomide is safe and effective for treating HHT patients, offering greater efficacy and fewer side effects compared to thalidomide^69^. These findings highlight the potential of repurposing cancer drugs as therapeutic options for HHT. In this study, we demonstrate that targeting the flow-mediated PIEZO1 signaling pathway could serve as a novel therapeutic strategy for HHT, offering new opportunities for effective intervention in HHT patients.

## Affiliations

Cardiovascular Research Center, Department of Internal Medicine, Yale University School of Medicine, New Haven CT, USA. (H.P, S.L, J.F, M.A.S, L.Y, A.E), Yale University School of Medicine, Department of Hematology (M.R.), Yale University School of Medicine, Departments of Cell Biology and Biomedical Engineering (M.A.S), Yale University School of Medicine, Department of Molecular and Cellular Physiology (A.E), Université de Paris, PARCC, INSERM, F-75006 Paris (A.E)

## Supplemental Figure legends

**Supplemental Figure 1. Validation of PIEZO1 expression in *Alk1* and *Piezo1* double ECKO.**(A) ALK1 (white), IB4 (magenta) and PIEZO1 (green) immunostaining of retinal flat mounts from P6 *Alk1^f/f^*, *Alk1^f/f^ Cdh5 Cre^ERT2^* and *Alk1^f/f^ Piezo1^f/f^ Cdh5 Cre^ERT2^* pups. (B) IB4 staining of retinal flat mounts from P6 *Alk1^f/f^*, *Alk1^f/f^ Cdh5 Cre^ERT2^*, *Alk1^f/f^ Piezo1^f/+^ Cdh5 Cre^ERT2^*, and *Alk1^f/f^ Piezo1^f/f^ Cdh5 Cre^ERT2^* mice. (C) Quantification of AVM number (n=10-12). One-way ANOVA with Sidak’s multiple comparisons test. (D) Quantification of AVM vessel diameter (n=10-12). One-way ANOVA with Sidak’s multiple comparisons test. (E) Kaplan-Meier survival curve, showing lethality from *Alk1^f/f^*, *Alk1^f/f^ Mfsd2a Cre^ERT2^*, *Alk1^f/f^ Piezo1^f/+^ Mfsd2a Cre^ERT2^* and *Alk1^f/f^ Piezo1^f/f^ Mfsd2a Cre^ERT2^* mice. Gene deletion was induced by intra-gastric injections with 100 μg TAM into pups at P4. TAM-injected *Cre^ERT2^* negative littermates were used as controls. Error bars: SEM. *P-value < 0.05, *** P-value < 0.001, ns: nonsignificant, Scale bars: 200 μm (A), 500 μm (E)

**Supplemental Figure 2. KLF4 upregulation in *Alk1* ECKO retinas and PIEZO1 inhibition reduces KLF4 expression.** (A) *Klf4* mRNA expression by qPCR in purified mouse retinas from P6 TAM-injected mice. n=5. Error bars: SEM. *P-value < 0.05, Mann–Whitney *U* test. (B) KLF4 (green) and IB4 (magenta) immunostaining of retinal flat mounts from P6 *Alk1^f/f^* and *Alk1^f/f^* Cdh5 *Cre^ERT2^* pups. (C) Immunostaining of KLF4 (white) and IB4 (magenta) in retinal flat mounts from P6 *Alk1^f/f^ Cdh5 Cre^ERT2^* mice injected with TAM at P4 and GsMTx4 or vehicle at P4 and P5. Scale bars: 500 μm (B and C)

## References

1. Shovlin CL. Hereditary haemorrhagic telangiectasia: Pathophysiology, diagnosis and treatment. Blood Reviews. 2010;24:203–219.

2. McDonald J, Bayrak-Toydemir P and Pyeritz RE. Hereditary hemorrhagic telangiectasia: An overview of diagnosis, management, and pathogenesis. Genetics in Medicine. 2011;13:607–616.

3. Govani FS and Shovlin CL. Hereditary haemorrhagic telangiectasia: A clinical and scientific review. European Journal of Human Genetics. 2009;17:860–871.

4. Bernabéu C, Blanco FJ, Langa C, Garrido-Martin EM and Botella LM. Involvement of the TGF-β superfamily signalling pathway in hereditary haemorrhagic telangiectasia. Journal of Applied Biomedicine. 2010;8:169–177.

5. Roman BL and Hinck AP. ALK1 signaling in development and disease: new paradigms. Cellular and Molecular Life Sciences. 2017;74:4539–4560.

6. Ruiz-Llorente L, Gallardo-Vara E, Rossi E, Smadja DM, Botella LM and Bernabeu C. Endoglin and alk1 as therapeutic targets for hereditary hemorrhagic telangiectasia. Expert Opinion on Therapeutic Targets. 2017;21:933–947.

7. McAllister KA, Grogg KM, Johnson DW, Gallione CJ, Baldwin MA, Jackson CE, Helmbold EA, Markel DS, McKinnon WC, Murrel J, McCormick MK, Pericak-Vance MA, Heutink P, Oostra BA, Haitjema T, Westerman CJJ, Porteous ME, Guttmacher AE, Letarte M and Marchuk DA. Endoglin, a TGF-β binding protein of endothelial cells, is the gene for hereditary haemorrhagic telangiectasia type 1. Nature Genetics. 1994;8:345–351.

8. Tual-Chalot S, Mahmoud M, Allinson KR, Redgrave RE, Zhai Z, Oh SP, Fruttiger M and Arthur HM. Endothelial depletion of Acvrl1 in mice leads to arteriovenous malformations associated with reduced endoglin expression. PLoS One. 2014;9:e98646.

9. Gallione CJ, Repetto GM, Legius E, Rustgi AK, Schelley SL, Tejpar S, Mitchell G, Drouin E, Westermann CJ and Marchuk DA. A combined syndrome of juvenile polyposis and hereditary haemorrhagic telangiectasia associated with mutations in MADH4 (SMAD4). Lancet. 2004;363:852–9.

10. Park H, Furtado J, Poulet M, Chung M, Yun S, Lee S, Sessa WC, Franco CA, Schwartz MA and Eichmann A. Defective Flow-Migration Coupling Causes Arteriovenous Malformations in Hereditary Hemorrhagic Telangiectasia. Circulation. 2021;144:805–822.

11. Singh E, Redgrave RE, Phillips HM and Arthur HM. Arterial endoglin does not protect against arteriovenous malformations. Angiogenesis. 2020;23:559–566.

12. Lee HW, Xu Y, He L, Choi W, Gonzalez D, Jin SW and Simons M. Role of Venous Endothelial Cells in Developmental and Pathologic Angiogenesis. Circulation. 2021;144:1308–1322.

13. Corti P, Young S, Chen CY, Patrick MJ, Rochon ER, Pekkan K and Roman BL. Interaction between alk1 and blood flow in the development of arteriovenous malformations. Development. 2011;138:1573–82.

14. Ola R, Dubrac A, Han J, Zhang F, Fang JS, Larrivée B, Lee M, Urarte AA, Kraehling JR, Genet G, Hirschi KK, Sessa WC, Canals FV, Graupera M, Yan M, Young LH, Oh PS and Eichmann A. PI3 kinase inhibition improves vascular malformations in mouse models of hereditary haemorrhagic telangiectasia. Nature Communications. 2016;7:13650.

15. Han C, Choe S-w, Kim YH, Acharya AP, Keselowsky BG, Sorg BS, Lee Y-J and Oh SP. VEGF neutralization can prevent and normalize arteriovenous malformations in an animal model for hereditary hemorrhagic telangiectasia 2. Angiogenesis. 2014;17:823–830.

16. Iriarte A, Figueras A, Cerdà P, Mora JM, Jucglà A, Penín R, Viñals F and Riera-Mestre A. PI3K (Phosphatidylinositol 3-Kinase) Activation and Endothelial Cell Proliferation in Patients with Hemorrhagic Hereditary Telangiectasia Type 1. Cells. 2019;8.

17. Jin Y, Muhl L, Burmakin M, Wang Y, Duchez AC, Betsholtz C, Arthur HM and Jakobsson L. Endoglin prevents vascular malformation by regulating flow-induced cell migration and specification through VEGFR2 signalling. Nature Cell Biology. 2017;19:639–652.

18. Alsina-Sanchís E, García-Ibáñez Y, Figueiredo AM, Riera-Domingo C, Figueras A, Matias-Guiu X, Casanovas O, Botella LM, Pujana MA, Riera-Mestre A, Graupera M and Viñals F. ALK1 Loss Results in Vascular Hyperplasia in Mice and Humans Through PI3K Activation. Arterioscler Thromb Vasc Biol. 2018;38:1216–1229.

19. Coon BG, Timalsina S, Astone M, Zhuang ZW, Fang J, Han J, Themen J, Chung M, Yang-Klingler YJ, Jain M, Hirschi KK, Yamamato A, Trudeau LE, Santoro M and Schwartz MA. A mitochondrial contribution to anti-inflammatory shear stress signaling in vascular endothelial cells. J Cell Biol. 2022;221.

20. Zheng Q, Zou Y, Teng P, Chen Z, Wu Y, Dai X, Li X, Hu Z, Wu S, Xu Y, Zou W, Song H and Ma L. Mechanosensitive Channel PIEZO1 Senses Shear Force to Induce KLF2/4 Expression via CaMKII/MEKK3/ERK5 Axis in Endothelial Cells. Cells. 2022;11.

21. Wang S, Chennupati R, Kaur H, Iring A, Wettschureck N and Offermanns S. Endothelial cation channel PIEZO1 controls blood pressure by mediating flow-induced ATP release. J Clin Invest. 2016;126:4527–4536.

22. Beech DJ and Kalli AC. Force Sensing by Piezo Channels in Cardiovascular Health and Disease. Arterioscler Thromb Vasc Biol. 2019;39:2228–2239.

23. Li J, Hou B, Tumova S, Muraki K, Bruns A, Ludlow MJ, Sedo A, Hyman AJ, McKeown L, Young RS, Yuldasheva NY, Majeed Y, Wilson LA, Rode B, Bailey MA, Kim HR, Fu Z, Carter DA, Bilton J, Imrie H, Ajuh P, Dear TN, Cubbon RM, Kearney MT, Prasad RK, Evans PC, Ainscough JF and Beech DJ. Piezo1 integration of vascular architecture with physiological force. Nature. 2014;515:279–282.

24. Ranade SS, Qiu Z, Woo SH, Hur SS, Murthy SE, Cahalan SM, Xu J, Mathur J, Bandell M, Coste B, Li YS, Chien S and Patapoutian A. Piezo1, a mechanically activated ion channel, is required for vascular development in mice. Proc Natl Acad Sci U S A. 2014;111:10347–52.

25. Park SO, Wankhede M, Lee YJ, Choi E-J, Fliess N, Choe S-W, Oh S-H, Walter G, Raizada MK, Sorg BS and Oh SP. Real-time imaging of de novo arteriovenous malformation in a mouse model of hereditary hemorrhagic telangiectasia. The Journal of Clinical Investigation. 2009;119:3487–3496.

26. Wang Y, Nakayama M, Pitulescu ME, Schmidt TS, Bochenek ML, Sakakibara A, Adams S, Davy A, Deutsch U, Lüthi U, Barberis A, Benjamin LE, Mäkinen T, Nobes CD and Adams RH. Ephrin-B2 controls VEGF-induced angiogenesis and lymphangiogenesis. Nature. 2010;465:483–6.

27. Pu W, Zhang H, Huang X, Tian X, He L, Wang Y, Zhang L, Liu Q, Li Y, Li Y, Zhao H, Liu K, Lu J, Zhou Y, Huang P, Nie Y, Yan Y, Hui L, Lui KO and Zhou B. Mfsd2a+ hepatocytes repopulate the liver during injury and regeneration. Nat Commun. 2016;7:13369.

28. Vanlandewijck M, He L, Mäe MA, Andrae J, Ando K, Del Gaudio F, Nahar K, Lebouvier T, Laviña B, Gouveia L, Sun Y, Raschperger E, Räsänen M, Zarb Y, Mochizuki N, Keller A, Lendahl U and Betsholtz C. A molecular atlas of cell types and zonation in the brain vasculature. Nature. 2018;554:475–480.

29. Zarkada G, Howard JP, Xiao X, Park H, Bizou M, Leclerc S, Künzel SE, Boisseau B, Li J, Cagnone G, Joyal JS, Andelfinger G, Eichmann A and Dubrac A. Specialized endothelial tip cells guide neuroretina vascularization and blood-retina-barrier formation. Dev Cell. 2021;56:2237–2251.e6.

30. Winkler EA, Kim CN, Ross JM, Garcia JH, Gil E, Oh I, Chen LQ, Wu D, Catapano JS, Raygor K, Narsinh K, Kim H, Weinsheimer S, Cooke DL, Walcott BP, Lawton MT, Gupta N, Zlokovic BV, Chang EF, Abla AA, Lim DA and Nowakowski TJ. A single-cell atlas of the normal and malformed human brain vasculature. Science. 2022;375:eabi7377.

31. Genet G, Genet N, Paila U, Cain SR, Cwiek A, Chavkin NW, Serbulea V, Figueras A, Cerdà P, McDonnell SP, Sankaranarayanan D, Huba M, Nelson EA, Riera-Mestre A and Hirschi KK. Induced Endothelial Cell Cycle Arrest Prevents Arteriovenous Malformations in Hereditary Hemorrhagic Telangiectasia. Circulation. 2024;149:944–962.

32. Banerjee K, Lin Y, Gahn J, Cordero J, Gupta P, Mohamed I, Graupera M, Dobreva G, Schwartz MA and Ola R. SMAD4 maintains the fluid shear stress set point to protect against arterial-venous malformations. J Clin Invest. 2023;133.

33. Ola R, Künzel Sandrine H, Zhang F, Genet G, Chakraborty R, Pibouin-Fragner L, Martin K, Sessa W, Dubrac A and Eichmann A. SMAD4 Prevents Flow Induced Arteriovenous Malformations by Inhibiting Casein Kinase 2. Circulation. 2018;138:2379–2394.

34. Dinakaran S, Qutaina S, Zhao H, Tang Y, Wang Z, Ruiz S, Nomura-Kitabayashi A, Metz CN, Arthur HM, Meadows SM, Blanc L, Faughnan ME and Marambaud P. CDK6-mediated endothelial cell cycle acceleration drives arteriovenous malformations in hereditary hemorrhagic telangiectasia. Nat Cardiovasc Res. 2024.

35. Levet S, Ciais D, Merdzhanova G, Mallet C, Zimmers TA, Lee SJ, Navarro FP, Texier I, Feige JJ, Bailly S and Vittet D. Bone morphogenetic protein 9 (BMP9) controls lymphatic vessel maturation and valve formation. Blood. 2013;122:598–607.

36. Olsen OE, Wader KF, Hella H, Mylin AK, Turesson I, Nesthus I, Waage A, Sundan A and Holien T. Activin A inhibits BMP-signaling by binding ACVR2A and ACVR2B. Cell Commun Signal. 2015;13:27.

37. Aspalter IM, Gordon E, Dubrac A, Ragab A, Narloch J, Vizán P, Geudens I, Collins RT, Franco CA, Abrahams CL, Thurston G, Fruttiger M, Rosewell I, Eichmann A and Gerhardt H. Alk1 and Alk5 inhibition by Nrp1 controls vascular sprouting downstream of Notch. Nat Commun. 2015;6:7264.

38. Wu M, Wu S, Chen W and Li YP. The roles and regulatory mechanisms of TGF-β and BMP signaling in bone and cartilage development, homeostasis and disease. Cell Res. 2024;34:101–123.

39. Cahalan SM, Lukacs V, Ranade SS, Chien S, Bandell M and Patapoutian A. Piezo1 links mechanical forces to red blood cell volume. Elife. 2015;4.

40. Wang J, Ma Y, Sachs F, Li J and Suchyna TM. GsMTx4-D is a cardioprotectant against myocardial infarction during ischemia and reperfusion. J Mol Cell Cardiol. 2016;98:83–94.

41. Ward CW, Sachs F, Bush ED and Suchyna TM. GsMTx4-D provides protection to the D2.mdx mouse. Neuromuscul Disord. 2018;28:868–877.

42. Suchyna TM. Piezo channels and GsMTx4: Two milestones in our understanding of excitatory mechanosensitive channels and their role in pathology. Prog Biophys Mol Biol. 2017;130:244–253.

43. Choi D, Park E, Yu RP, Cooper MN, Cho IT, Choi J, Yu J, Zhao L, Yum JI, Yu JS, Nakashima B, Lee S, Seong YJ, Jiao W, Koh CJ, Baluk P, McDonald DM, Saraswathy S, Lee JY, Jeon NL, Zhang Z, Huang AS, Zhou B, Wong AK and Hong YK. Piezo1-Regulated Mechanotransduction Controls Flow-Activated Lymphatic Expansion. Circ Res. 2022;131:e2–e21.

44. Anzell AR, Kunz AB, Donovan JP, Tran TG, Lu X, Young S and Roman BL. Blood flow regulates acvrl1 transcription via ligand-dependent Alk1 activity. Angiogenesis. 2024;27:501–522.

45. dela Paz NG, Melchior B and Frangos JA. Early VEGFR2 activation in response to flow is VEGF-dependent and mediated by MMP activity. Biochem Biophys Res Commun. 2013;434:641–6.

46. dela Paz NG, Walshe TE, Leach LL, Saint-Geniez M and D’Amore PA. Role of shear-stress-induced VEGF expression in endothelial cell survival. J Cell Sci. 2012;125:831–43.

47. Clark PR, Jensen TJ, Kluger MS, Morelock M, Hanidu A, Qi Z, Tatake RJ and Pober JS. MEK5 is activated by shear stress, activates ERK5 and induces KLF4 to modulate TNF responses in human dermal microvascular endothelial cells. Microcirculation. 2011;18:102–17.

48. Buschbeck M and Ullrich A. The unique C-terminal tail of the mitogen-activated protein kinase ERK5 regulates its activation and nuclear shuttling. J Biol Chem. 2005;280:2659–67.

49. Morimoto H, Kondoh K, Nishimoto S, Terasawa K and Nishida E. Activation of a C-terminal transcriptional activation domain of ERK5 by autophosphorylation. J Biol Chem. 2007;282:35449–56.

50. Mody N, Leitch J, Armstrong C, Dixon J and Cohen P. Effects of MAP kinase cascade inhibitors on the MKK5/ERK5 pathway. FEBS Lett. 2001;502:21–4.

51. Ohnesorge N, Viemann D, Schmidt N, Czymai T, Spiering D, Schmolke M, Ludwig S, Roth J, Goebeler M and Schmidt M. Erk5 activation elicits a vasoprotective endothelial phenotype via induction of Kruppel-like factor 4 (KLF4). J Biol Chem. 2010;285:26199–210.

52. Varia MA, Calkins-Adams DP, Rinker LH, Kennedy AS, Novotny DB, Fowler WC, Jr. and Raleigh JA. Pimonidazole: a novel hypoxia marker for complementary study of tumor hypoxia and cell proliferation in cervical carcinoma. Gynecol Oncol. 1998;71:270–7.

53. Suchting S, Freitas C, le Noble F, Benedito R, Bréant C, Duarte A and Eichmann A. The Notch ligand Delta-like 4 negatively regulates endothelial tip cell formation and vessel branching. Proc Natl Acad Sci U S A. 2007;104:3225–30.

54. Li X, Kordsmeier J, Nookaew I, Kim HN and Xiong J. Piezo1 stimulates mitochondrial function via cAMP signaling. Faseb j. 2022;36:e22519.

55. Coon BA-O, Timalsina SA-O, Astone MA-O, Zhuang ZA-O, Fang JA-O, Han J, Themen J, Chung MA-O, Yang-Klingler YA-O, Jain M, Hirschi KK, Yamamato AA-O, Trudeau LA-O, Santoro MA-O and Schwartz MA-O. A mitochondrial contribution to anti-inflammatory shear stress signaling in vascular endothelial cells. LID - 10.1083/jcb.202109144 [doi] LID - e202109144.

56. Tusa I, Menconi A, Tubita A and Rovida E. Pathophysiological Impact of the MEK5/ERK5 Pathway in Oxidative Stress. Cells. 2023;12.

57. Verhoeven J, Baelen J, Agrawal M and Agostinis P. Endothelial cell autophagy in homeostasis and cancer. FEBS Lett. 2021;595:1497–1511.

58. Schaaf MB, Houbaert D, Meçe O and Agostinis P. Autophagy in endothelial cells and tumor angiogenesis. Cell Death Differ. 2019;26:665–679.

59. Ng I, Tan WL, Ng PY and Lim J. Hypoxia inducible factor-1alpha and expression of vascular endothelial growth factor and its receptors in cerebral arteriovenous malformations. J Clin Neurosci. 2005;12:794–9.

60. Wang L, Guo S, Zhang N, Tao Y, Zhang H, Qi T, Liang F and Huang Z. The role of SDF-1/CXCR4 in the vasculogenesis and remodeling of cerebral arteriovenous malformation. Ther Clin Risk Manag. 2015;11:1337–44.

61. Meijer-Jorna LB, Aronica E, van der Loos CM, Troost D and van der Wal AC. Congenital vascular malformations--cerebral lesions differ from extracranial lesions by their immune expression of the glucose transporter protein GLUT1. Clin Neuropathol. 2012;31:135–41.

62. Wang Z, Chen J, Babicheva A, Jain PP, Rodriguez M, Ayon RJ, Ravellette KS, Wu L, Balistrieri F, Tang H, Wu X, Zhao T, Black SM, Desai AA, Garcia JGN, Sun X, Shyy JY, Valdez-Jasso D, Thistlethwaite PA, Makino A, Wang J and Yuan JX. Endothelial upregulation of mechanosensitive channel Piezo1 in pulmonary hypertension. Am J Physiol Cell Physiol. 2021;321:C1010–c1027.

63. Cerdà P, Castillo SD, Aguilera C, Iriarte A, Rocamora JL, Larrinaga AM, Viñals F, Graupera M and Riera-Mestre A. New genetic drivers in hemorrhagic hereditary telangiectasia. Eur J Intern Med. 2024;119:99–108.

64. Bernabeu CA-O, Bayrak-Toydemir P, McDonald J and Letarte M. Potential Second-Hits in Hereditary Hemorrhagic Telangiectasia. LID - 10.3390/jcm9113571 [doi] LID - 3571.

65. Al Tabosh T, Al Tarrass M, Tourvieilhe L, Guilhem A, Dupuis-Girod S and Bailly S. Hereditary hemorrhagic telangiectasia: from signaling insights to therapeutic advances. J Clin Invest. 2024;134.

66. Ardelean DS and Letarte M. Anti-angiogenic therapeutic strategies in hereditary hemorrhagic telangiectasia. Front Genet. 2015;6:35.

67. Droege F, Thangavelu K, Lang S and Geisthoff U. Improvement in hereditary hemorrhagic telangiectasia after treatment with the multi-kinase inhibitor Sunitinib. Ann Hematol. 2016;95:2077–2078.

68. Geisthoff UW, Nguyen HL and Hess D. Improvement in hereditary hemorrhagic telangiectasia after treatment with the phosphoinositide 3-kinase inhibitor BKM120. Ann Hematol. 2014;93:703–4.

69. Al-Samkari H, Kasthuri RS, Iyer VN, Pishko AM, Decker JE, Weiss CR, Whitehead KJ, Conrad MB, Zumberg MS, Zhou JY, Parambil J, Marsh D, Clancy M, Bradley L, Wisniewski L, Carper BA, Thomas SM and McCrae KR. Pomalidomide for Epistaxis in Hereditary Hemorrhagic Telangiectasia. N Engl J Med. 2024;391:1015–1027.

