## Supplementary figures and images for "PIEZO1 overexpression in hereditary hemorrhagic telangiectasia arteriovenous malformations"

### Supplementary Figures

Supplementary Fig 1

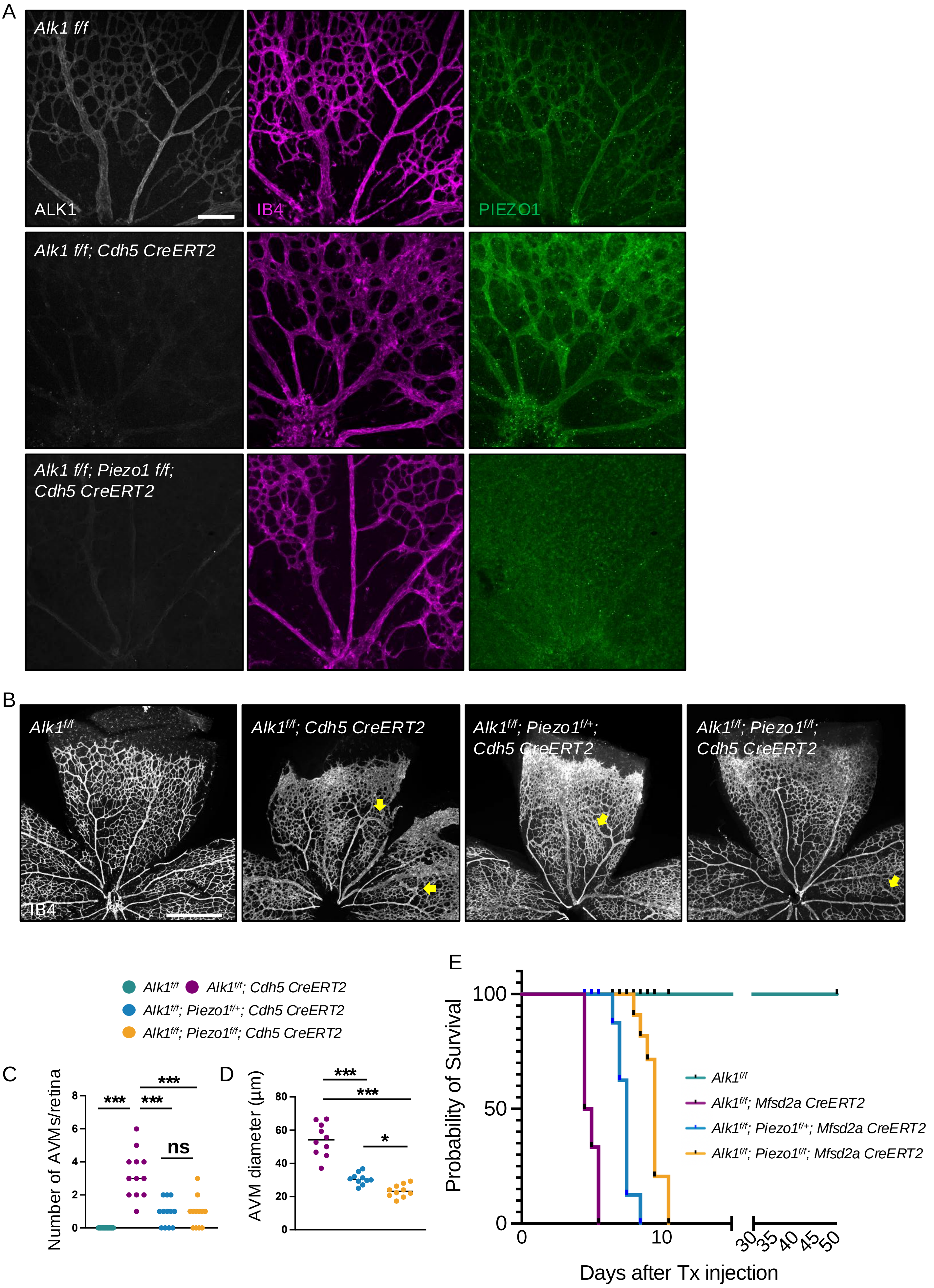

Supplementary Fig 2

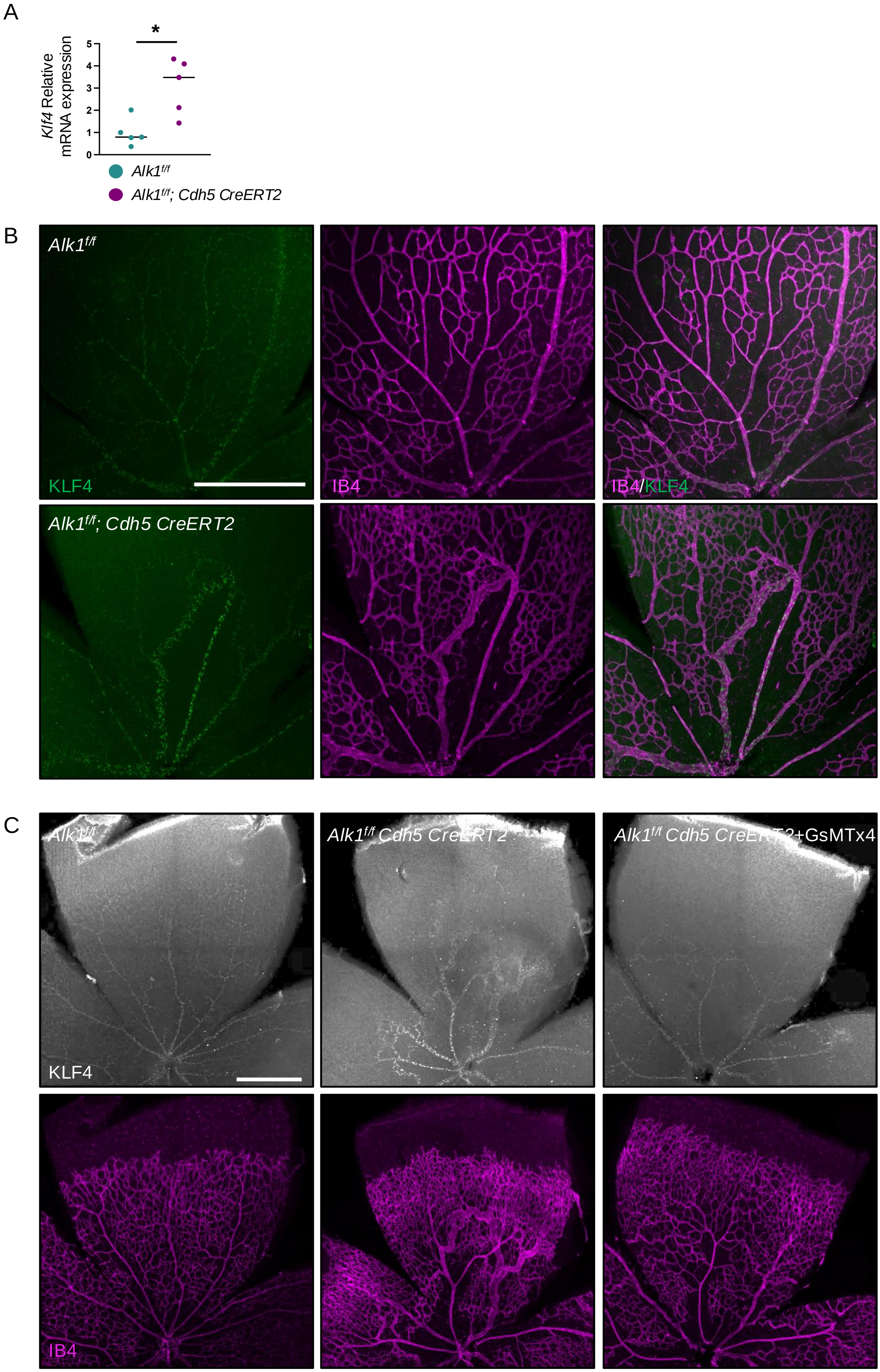
